# Transcription promotes the restoration of chromatin following DNA replication

**DOI:** 10.1101/2023.04.19.537523

**Authors:** Susanne Bandau, Vanesa Alvarez, Hao Jiang, Sarah Graff, Ramasubramanian Sundaramoorthy, Matt Toman, Tom Owen-Hughes, Simone Sidoli, Angus Lamond, Constance Alabert

## Abstract

DNA replication results in the transient eviction of nucleosomes, RNAPII and transcription regulators. How chromatin organization is duplicated on the two daughter strands is a central question in epigenetics. In mammals, transcription restarts on newly replicated DNA within a couple of hours, promoting chromatin accessibility. However, the role of transcription in the restoration of other chromatin determinants following DNA replication remains unclear. Here we have monitored protein re-association to newly replicated DNA upon inhibition of transcription using iPOND coupled to quantitative mass spectrometry. We show that nucleosome assembly and the re-establishment of most histone modifications are uncoupled from transcription restart. However, upon transcription inhibition, the re-association of many proteins was altered, including ATP-dependent remodellers, transcription regulators, the histone variant H2A.Z, histone modifiers as well as the restoration of H3.3K36me2. Finally, transcription also provoked the recruitment of several DNA repair proteins, revealing that transcription promotes chromatin reestablishment post-replication but is also a potential source of genotoxic stress.

## INTRODUCTION

Chromatin structures contain multiple layers of information that sustain transcription programs. These structures must be plastic enough to permit the establishment of distinct transcription programs in distinct cells during development, and robust enough to ensure the maintenance of established programs in proliferating cells. In cycling cells chromatin is profoundly modified twice. In mitosis, chromatin undergoes a rapid cycle of compaction and decompaction, allowing the distribution of the copied genetic material between the two daughter cells. In S phase, at each site of DNA synthesis, chromatin undergoes a cycle of disruption ahead of the replisome and of reassembly on the two daughter strands. As the genome is replicated following a temporal order, disruption of chromatin structures is genome wide but not simultaneous. Yet, any given time, ∼5000 forks progress on chromatin, evicting thousands of nucleosomes per minute. Despite major recent advances in our understanding of this dynamic process, the mechanisms that ensure the reassembly of chromatin structures on newly replicated DNA and their coordination with other DNA based processes remain unclear. Understanding how chromatin-based information propagate from cell to cell, and how this information is altered in a multitude of disease contexts such as cancers, is a central question in biology.

Following DNA replication, nucleosomes are first assembled with less well-defined positioning and transcription factor (TF) occupancy at gene regulatory sequences is reduced (Owens et al., 2019; Ramachandran and Henikoff, 2016; Stewart-Morgan et al., 2019). RNA polymerase II (RNAPII) is recruited to replicated DNA and the initiation competent form, RNAPII phosphorylated on serine 5 (RNAPII-pS5), progressively increases from 30 min to 2 hours following the passage of the replisome (Stewart-Morgan et al., 2019). Although there is no direct measure of *de novo* transcriptional activity on newly replicated chromatin, it is believed that similar to mitosis (Palozola et al., 2017), transcription is transiently reduced behind replisomes. Within 2 hours, pre-replicated RNAPII-pS5 levels and DNA accessibility are re-established (Stewart-Morgan et al., 2019). This restoration is not homogenous, as regions such as super-enhancers and CTCF binding sites are restored prior to promoters (Owens et al., 2019; Stewart-Morgan et al., 2019). Mechanistically, recent work has shown that in mouse ES cells, transcription is driving nucleosome re-positioning within gene regulatory regions (Stewart-Morgan et al., 2019). Yet, how transcription promotes nucleosome positioning remains unclear. Several chromatin remodelling enzymes, that have the ability to reposition or evict nucleosomes, are present on newly replicated DNA (Alvarez et al., 2023; Ramachandran and Henikoff, 2016). However, TFs defined as pioneers bind with similar kinetics as other TFs (Alvarez et al., 2023). Therefore, whether the most widely accepted model of pioneer TFs recruited first, followed by chromatin remodellers and RNAPII applies behind replisomes remains to be established.

Beyond nucleosome positioning, transcription promotes other processes taking place on newly replicated DNA such as the restoration of histone modifications. Following DNA replication, many histone methylations are diluted by the addition of newly synthesized histones that in mammals are largely unmethylated (Alabert et al., 2015; Reveron-Gomez et al., 2018). Several mechanisms have been proposed to overcome this dilution, such as the direct recruitment of H3K9me3 writers to the replisome (Padeken et al., 2022). H3K4me3 restoration following DNA replication has been suggested to rely on transcription (Reveron-Gomez et al., 2018; Serra-Cardona et al., 2022). A probable other candidate is H3K36me3, as its writer NSD2 directly bind RNAPII-pS2-S5, and this interaction promotes H3K36me3 deposition on chromatin (Neri et al., 2017; Venkatesh et al., 2012). More recently, H2BK120ub1 has been shown to be coupled with transcription (Flury et al., 2023). Yet, it remains unknown whether transcription is required for the restoration of other histone modifications. More generally, it is uncertain whether the mechanisms that maintain histone modifications in steady state chromatin are applicable on newly replicated chromatin. These questions have gained a novel interest following development of inhibitors and rapid degradation-based approaches. These studies revealed that continuous action of chromatin remodelling enzymes is required to maintain chromatin accessibility at enhancers (Blumli et al., 2021; Iurlaro et al., 2021; Schick et al., 2021). Similarly, the continuous presence of PRC1 at Polycomb target genes is required to maintain silencing (Dobrinic et al., 2021). Therefore, the finding that chromatin states are much more dynamic than anticipated further highlights the need to better understand the immediate response to chromatin disruption.

Here we examined the contribution of transcription to restoration of chromatin states on newly replicated DNA. To do so we use inhibitors to block RNAPII binding and elongation and analysed protein binding kinetics and restoration of histone modification behind replisomes using isolation of Proteins On Newly replicated DNA (iPOND) coupled to mass spectrometry (Sirbu et al., 2012). We show that the binding of RNAPII promotes the recruitment of all the ATP-dependent chromatin remodellers and of several TFs to newly replicated DNA. RNAPII elongation further ensures the retention on replicated chromatin of the BAF complex, INO80/SWR and otherwise steady TFs. Moreover, transcription promotes the re-assembly of H2A.Z and the restoration of H3.3K36me2, while simultaneously provoking the recruitment of several DNA repair factors. Finally, blocking transcription leads to the accumulation of the linker histone H1.X and perturbs the incorporation of H3K9me3 and H4K20me2. Altogether this study provides a comprehensive proteomics analysis assessing the effect of transcription inhibition on protein binding kinetics and histone modification restoration on post-replicated chromatin. It revealed that transcription promotes chromatin reestablishment post-replication but is also a potential source of genotoxic stress.

## RESULTS

### Isolation of newly replicated chromatin under conditions where transcription is inhibited

To examine the role of transcription in the restoration of chromatin following DNA replication (Fig. 1A), primary like foetal human lung fibroblasts (TIG-3) were used, and newly replicated chromatin was examined in the presence of transcription inhibitors using iPOND coupled to TMT mass spectrometry. Two small molecule inhibitors targeting two different transcription steps were selected. Triptolide (TPL) inhibits the ATPase activity of XPB, a subunit of TFIIH, which is part of the pre-initiation complex. This drug induces the rapid proteasomal degradation of RBP1 and therefore blocks RNAPII loading (Vispe et al., 2009). 5,6-dichlorobenzimidazone-1-β-D-ribofuranoside (DRB) inhibits the kinase activity of CDK9, a component of the transcription elongation factor pTEFb, thereby initially increasing RNAPII pausing followed by a reduction of the fraction of chromatin bound RNAPII (Steurer et al., 2018). To establish the drug treatment conditions required to impair transcription in TIG-3 cells, we used Quantitative Image Based Cytometry (QIBC) (Toledo et al., 2013). Within 2 hours, 1 μM of TPL and 50 μM of DRB significantly reduced transcription rates (Fig. 1B, S1A). TPL treatment also reduced the level of chromatin bound RNAPII (Fig. 1C), which is consistent with the previously described rapid induction of RNAPII proteasomal degradation. Importantly, in these conditions DNA replication was unaffected (Fig. 1D, 1E), making it possible to label newly replicated DNA in conditions where transcription is impaired.

**Figure 1:**
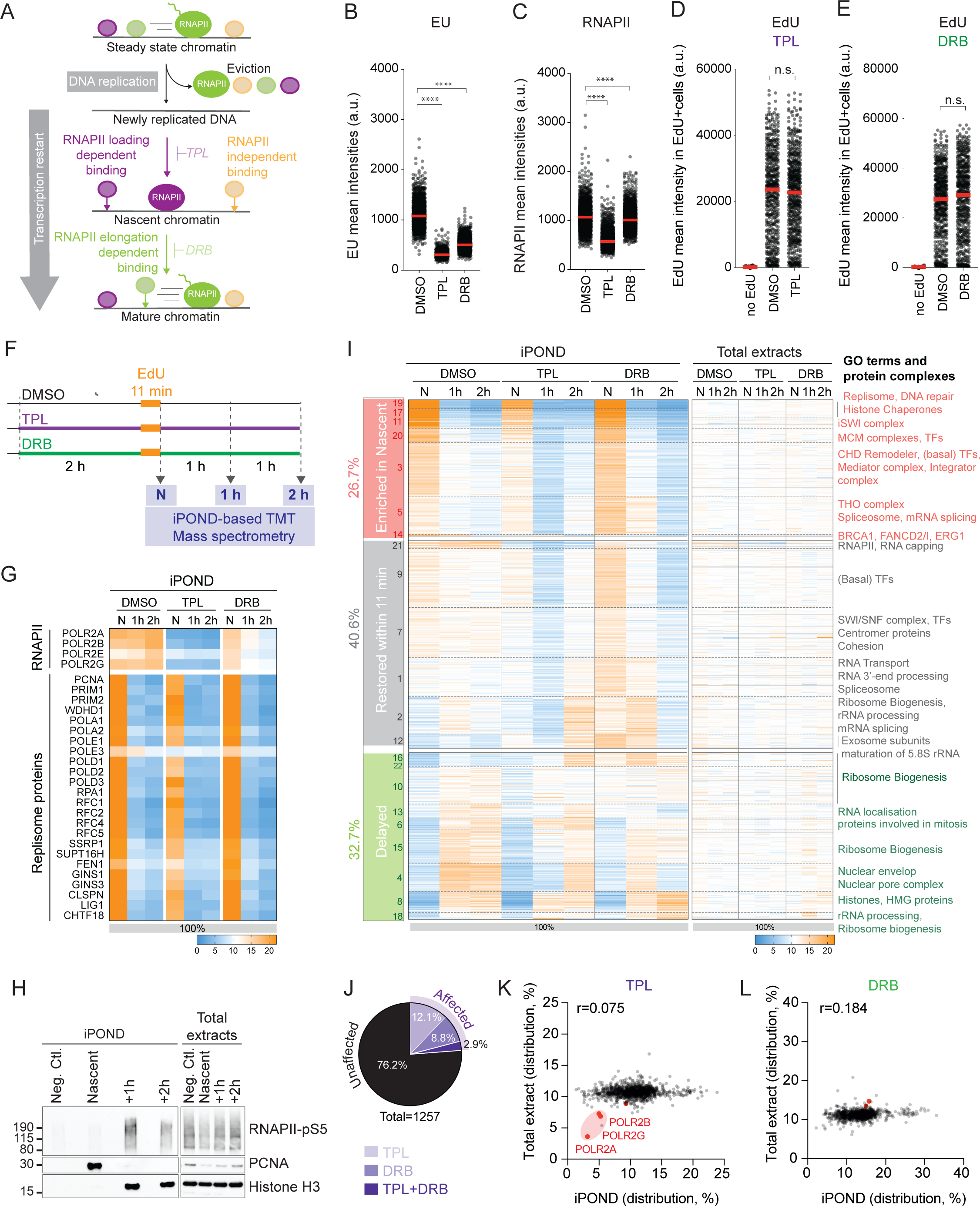
Proteomic profiling of chromatin behind replisomes upon transcription inhibition. **A.** Behind replisomes, histones and chromatin components are reassembled onto newly replicated DNA. How transcription contributes to this process is unclear. **B-E.** QIBC based analysis of EU level (C), chromatin bound RNAPII (D), and EdU level (E, F) in DMSO, TPL and DRB treated cells. Graphs show the mean intensity per nuclei, > 539 nuclei were analysed per sample. Red line: median; p value was calculated using an unpaired t test. **F**. Experimental design of the iPOND-TMT time course experiment in TIG-3 cells in the presence of DMSO, TPL or DRB. **G**. Heatmap of relative abundance of identified replisome components and RNAPII subunits shown as percentage of enrichment (n = 3). Each column represents a time point (N: Nascent, 1h, and 2h) and each row corresponds to the protein indicated on the left. The sum of each row corresponds to 100% of enrichment. Colour scale indicated and used throughout Figure 1. **H**. Western Blot analysis of iPOND samples. The Western Blot was probed with antibodies against RNAPII-pS5, PCNA, and Histone H3 (indicated on the right). Samples label shown on top. **I**. Hierarchical clustering using 22 clusters (1257 proteins, n = 3). Cluster numbers are indicated on the left and in Table S1. Clusters are grouped into three groups: Enriched in nascent (red), Restored within 11 min (grey), and Delayed (green). The number of proteins in each group is shown as percentage on the left. For each protein identified, the percentage of enrichment from total cell extracts is aligned on the right of the clustered iPOND. GO terms and protein groups are shown on the right. See Figure S1F for TPL and DRB effects on each category. **J**. Pie chart summarising the effect of TPL and DRB treatment compared to DMSO. The proteins were divided into four groups: Unaffected (black), Affected (purple), Affected by TPL only and Affected by DRB only (light purples). **K, L**. Scatter plots comparing the relative percentage of enrichment between nascent iPOND samples and the respective cellular extracts in TPL (K) and DRB (L) treated cells. The Pearson correlation coefficient (r) is shown.

Newly replicated DNA was labelled using an 11-minute pulse of EdU, covering the entire genome as the cell population was left asynchronous (Reveron-Gomez et al., 2018) (Fig. 1F). At the end of the EdU pulse, cells were either processed according to the iPOND protocol (Nascent sample) or chased using thymidine for an additional 1 hour (+1h sample) or 2 hours (+2h sample). The thymidine chase stops EdU incorporation and therefore enables the fate of EdU-labelled DNA to be monitored (Sirbu et al., 2011). Three independent biological replicates were performed (Fig. S1B, S1C) and 4933 proteins identified, of which 1257 proteins were identified in all conditions (Table S1). This dataset was used to calculate the relative abundance of each identified proteins on labelled DNA, thereby revealing their binding kinetics following DNA replication.

The relative abundances between time points were calculated for each protein and expressed as percentage in a heatmap format (Fig. 1G). Core replisome components were enriched at the nascent time point compared to later time points in DMSO, TPL and DRB, indicating that nascent and mature chromatin has been successfully isolated in all conditions. In DMSO, RNAPII abundance only moderately increased between 11 and 120 minutes after the passage of replisomes (Fig. 1G, S1D). To assess the status of RNAPII we repeated the experiment and probed for RNAPII serine 5 phosphorylation (RNAPII-pS5), a phosphorylation deposited on RNAPII upon its activation. Consistent with CHOR-seq in mES (Stewart-Morgan et al., 2019), RNAPII-pS5 levels were low in the 11 minutes sample (nascent) compared to 2 hours after the passage of the fork (Fig. 1H, S1D). Together with the mass spectrometry results, this confirms that behind replisomes RNAPII binds within the first 11 minutes and is activated within the following hour. Upon TPL treatment, the abundance of RNAPII components was greatly reduced on replicated chromatin compared to DMSO, indicating that RNAPII binding has been successfully impaired (Fig. 1G). Upon DRB treatment, consistent with *in vitro* FRAP experiments (Steurer et al., 2018), RNAPII peaked on nascent chromatin before decreasing. Therefore, newly replicated chromatin has been successfully isolated in conditions where RNAPII binding and elongation was permitted (DMSO), RNAPII binding was prevented (TPL), or RNAPII binding had been allowed but transcription elongation prevented (DRB). Of note, RNAPI and RNAPIII levels were affected by TPL and DRB, but effects were dampened compared to RNAPII (Fig. S1E).

### RNAPII’s binding and elongation contribute to protein binding to newly replicated DNA

To capture the different phases of chromatin restoration, hierarchical clustering was performed on the data set based on relative protein abundance across the 3 time points and 3 conditions (Fig. 1I). Using the Nascent time point as the starting point of chromatin restoration (the first 11 minutes), and +2h as the end point, clusters were grouped into three categories: clusters showing a 1.2-fold or more enrichment between the Nascent time point and any other time point were named “Enriched in nascent”. Clusters showing a 1.2 to 0.83-fold enrichment between Nascent and +2h were named “Restored within 11 min”, as they included proteins equally abundant 11 minutes after the passage of the fork and 2 hours later. Finally, clusters showing a 0.83-fold or less enrichment between Nascent and +2h were named “Delayed”. Consistent with previous work (Alvarez et al., 2023), clustering analysis indicates that in DMSO, most chromatin components are restored within the first 11 minutes after the passage of the replication fork, while for 32.7% of the proteins identified, chromatin restoration is not as prompt (Fig. 1I, S1F).

To identify the proteins whose binding on replicated chromatin was the most affected by blocking transcription, we compared the composition of newly replicated chromatin in DMSO to TPL and DRB treated cells. The binding of 24% of the proteins identified were significantly affected by transcription inhibition, TPL having the strongest effect (Fig. 1J). Treating cells with TPL overall delayed the recruitment of proteins especially at the 2-hr time point compared to DMSO conditions as well as causing the accumulation others (Fig. 1I, S1F). This includes proteins involved in mRNA processing, splicing and ribosome biogenesis. In contrast, treating cells with DRB accelerated the recruitment of many proteins resulting in increased recruitment to nascent chromatin (Fig1I, Fig S1F). The cohesin complex provides an example of the overall effect of TPL and DRB treatments (Fig. S1G, S1H). Taken together this analysis revealed that blocking transcription perturbs the restoration of newly replicated chromatin.

As RNA production was purposely blocked using transcription inhibitors, we first tested whether the changes in binding kinetics observed upon DRB or TPL treatment could be explained by a decrease in abundance of these proteins in the cell. To do so, total cellular extracts were analysed by TMT quantitative mass spectrometry in parallel with each iPOND sample (Table S1). As expected from previous studies (Vispe et al., 2009), the abundance of several proteins decreased upon TPL treatment (Fig. S1I). Yet, the changes of abundance detected on replicated chromatin upon TPL or DRB treatments could not be inferred from the changes in abundance observed in the cells (Fig. 1I, right: cellular abundance aligned to iPOND clustering). Moreover, the composition of total cell extracts and replicated chromatin fractions showed very low to no positive Pearson’s correlations (Fig. 1K, 1L, S1J), the exception being RNAPII itself in TPL treated cells as expected (Fig. 1K). As blocking transcription for a couple of hours may primarily affect the abundance of short half-life proteins and low abundant proteins, we selected the shortest half-life proteins (Li et al., 2021), and used iBAQ as a proxy to select the least abundant proteins, and performed the same analysis as in Figure 1J. The lowest abundant proteins, as well as proteins with the shortest half-life were not more affected by TPL or DRB treatment (Fig. S1K). Altogether these results reveal that transcription inhibition affects the binding kinetics of hundreds of proteins, and this is not an indirect consequence of changes in protein abundance.

### Blocking transcription reduced the recruitment of DNA repair proteins

Detailed inspection of nascent chromatin composition revealed that upon DRB and TPL treatments, newly replicated chromatin was depleted in proteins involved in the regulation of transcription such as chromatin remodellers, transcription regulators, and histone modifiers, as well as DNA repair proteins (Fig. 2A-C, S2A-D). Importantly, and as shown for the clusters defined in Figure 1, these depletions were not an indirect consequence of blocking protein production (Fig. S2E, S2F). Moreover, only 4/50 of the proteins whose binding is reduced upon TPL treatment are known RNAPII interactors (Ebmeier et al., 2017) (Fig. S2G). This indicates that our data captures direct and indirect dependencies on RNAPII, and that most of the identified RNAPII interactors are loaded onto nascent chromatin independently of RNAPII.

**Figure 2:**
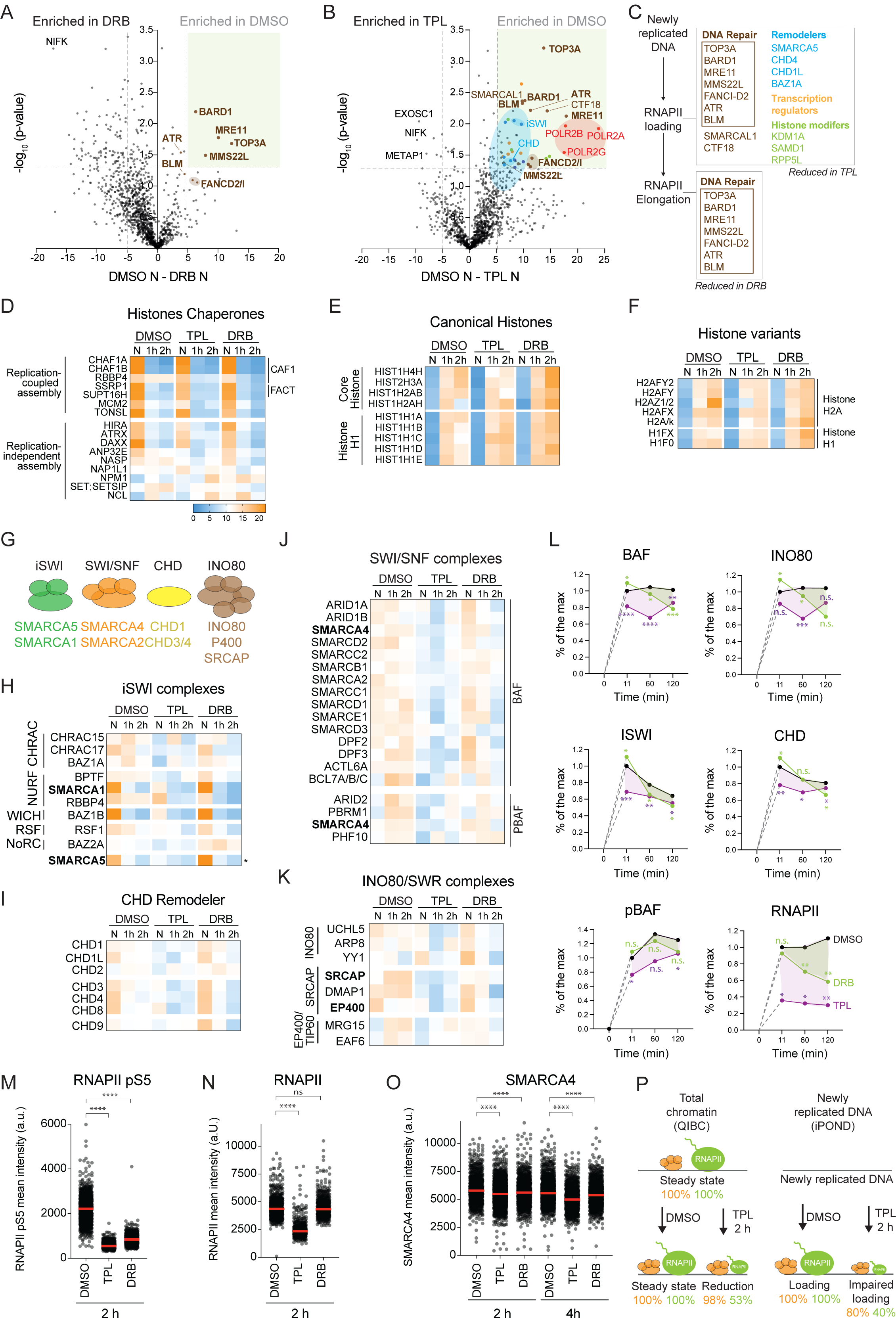
Chromatin remodellers abundance on nascent and steady state chromatin is impaired upon TPL treatment. **A, B**. Enrichment of proteins across DRB and TPL iPOND-TMT-MS datasets. RNAPII, red; DNA repair, brown; Chromatin remodellers, blue; Transcription regulators, orange; Histone modifiers, green. **C.** Protein families affected by TPL and DRB treatments. **D-F.** Heatmaps of relative enrichment of Histone Chaperones (D), Canonical Histones (E), and Histone variants (F) (n = 3). Each column represents a time point (N: Nascent, 1h, and 2h) and each row corresponds to the protein indicated on the left. The sum of each row corresponds to 100% of enrichment. Colour scale is indicated. **G.** The four families of ATP-dependent chromatin remodelling complexes and their catalytic subunits below (ATPases). **H-K.** Same as in D-F for chromatin remodellers classified by families: iSWI (H), CHD (I), SWI/SNF (J), and INO80/SWR (K). **L.** The average of the relative enrichment for each remodeller family, between DMSO (black), TPL (purple), and DRB (green) treated cells. Standard deviation is shown, and p value calculated using a paired t test. **M-O**. QIBC analysis of chromatin bound intensities of RNAPII-pS5, RNAPII and SMARCA4 in DMSO, TPL or DRB treated cells. Graphs show the mean intensity per nuclei, >291 nuclei were analysed per sample. Red line: median; p value calculated using an unpaired t test. N=3, one representative experiment is shown. **P.** Effect of TPL treatment on RNAPII (green) and SMARCA4 (orange) in nascent chromatin and total chromatin.

As transcription is known to be a major source of replication stress (Berti et al., 2020), the reduced binding of several DNA repair proteins was expected, yet only certain pathways were involved (Fig. S2H-J). Upon DRB treatment, TOP3A, BLM, BARD11, MRE11, ATR, MMS22L, FANCD2, and FANCI were less abundant on nascent chromatin (Fig. 2A, brown). TOP3A and BLM promotes DNA hemicatenate resolution and are involved in several DNA repair pathways (Pommier et al., 2022). The other proteins are DNA repair proteins involved in fork quality control and HR as well as transcription replication conflicts (Berti et al., 2020). Upon TPL treatment, the binding of two more DNA repair proteins were affected, SMARCAL1 and CTF18 (Fig. 2B, brown). As these two proteins are protecting stalled replisomes, it suggests that RNAPII binding on newly replicated DNA may affect replisome progression. Altogether, this analysis further confirms that RNAPII’s binding and elongation on newly replicated chromatin are a source of genotoxic stress, and identifies dedicated repair factors handling transcription replication conflicts.

### RNAPII stabilizes association of many chromatin remodelling enzymes on newly replicated chromatin

The second group of proteins whose abundance was reduced in TPL treated cells were the chromatin remodellers (Fig. 2B, 2C). This result was intriguing as it suggests that behind replisomes, RNAPII loading may facilitate the association of these enzymes. It prompted us to examine in further detail the assembly of nucleosomes and the association of transcriptional regulators behind replisomes. As expected, replication coupled histone chaperones were enriched on nascent chromatin (Fig. 2D) (Alvarez et al., 2023). Association of canonical histones (H3, H4, H2A and H2B) was partial at 11 minutes and increased over the following hours (Fig. 2E). Together with chromatin accessibility studies (Ramachandran and Henikoff, 2016; Stewart-Morgan et al., 2019), this result reveals that within the first 11 minutes after the passage of the fork, newly assembled nucleosomes are poorly positioned and less abundant. Finally, the linker histone H1, and the histone variants detected, were also assembled within the first hour after the passage of the replisome (Fig. 2E, 2F).

Upon inhibition of transcription, association of histone chaperones and canonical histones was generally not strongly affected (Fig. 2D, 2E). The two exceptions were the histone variants H2A.Z (.1 and .2) and H1.X (H1FX) (Fig. 2F). H2A.Z deposition is known to be tightly coupled to transcription (Flury et al., 2023; Giaimo et al., 2019), and the finding that H2A.Z abundance was lower in DRB and TPL treated cells supports a role for transcription in the localisation of H2A.Z on replicated DNA. On the other hand, H1.X deposition is known to be replication uncoupled but its function remains unclear (Millan-Arino et al., 2016). As H1.X accumulates to significantly higher levels on replicated chromatin in DRB treated cells, it suggests that H1.X association to replicated chromatin is affected by a process linked to RNAPII-S5P.

The four main families of chromatin remodellers were also identified, SWI/SNF (BAF and pBAF), iSWI, CHD and INO80/SWR (Fig. 2G), and could be divided into three groups based on their binding kinetics in DMSO. iSWI and most CHD complexes were transiently enriched on newly replicated chromatin, BAF and INO80/SWR complexes remained stably bound to replicated chromatin, and the pBAF complex gradually increased (Fig. 2H-L). Upon TPL treatment, a 20% reduction in the association of most remodellers is observed. The 20% reduction in occupancy is statistically significant for all enzyme classes, but smaller in scale than the 60% reduction in RNAPII association (Fig. 2L). As a control, as TPL blocks the ATPase activity of XPB, we tested whether TPL could be blocking the ATPase activity of the remodelling enzymes, leading to changes in their association with chromatin (Kim et al., 2021). To do so, we selected SMARCA4 (BRG1) and examined its ability to slide nucleosomes upon TPL treatment *in vitro*. In this assay, TPL treatment did not impair the ability of SMARCA4 to slide nucleosomes (Fig. S2K). Taken together, these results indicate a role for RNAPII in stabilising the association of remodellers with newly replicated chromatin. This stabilisation is likely to involve RNAPII-S5P as enzymes other than pBAF show transiently enhanced recruitment to nascent chromatin reminiscent of the association of RNAPII following DRB treatment (Fig. 2L).

To test if the reduction in RNAPII and dissociation of chromatin remodellers from newly replicated chromatin upon TPL treatment was specific to newly replicated chromatin or could be applicable to total chromatin, we measured the level of RNAPII and SMARCA4 on total chromatin after treating cells with TPL. To do so, cells were pre-extracted with 0.5% triton prior to fixation, and chromatin bound RNAPII and SMARCA4 levels measured by QIBC. As controls, EU labelling and RNAPII-pS5 were included in the analysis. Upon TPL treatment, EU intensities and chromatin bound RNAPII-pS5 levels decreased (Fig. 2M, S2L). RNAPII as well as SMARCA4 abundance on chromatin were low in TPL compared to the DMSO condition (Fig. 2N, 2O). However, the loss of SMARCA4 on total chromatin was modest compared to nascent chromatin (after a 2 hr TPL treatment, 2% reduction on total chromatin, 20% reduction on nascent chromatin, Fig. 2L, 2O, 2P). Taken together, these results indicate that RNAPII mediated stabilisation of SMARCA4 is greater in nascent chromatin than in total chromatin.

### TF access to newly replicated DNA is stimulated by RNAPII binding and elongation

The third group of proteins affected by transcription inhibition are transcription regulators. In total, 98 TFs were identified in the TMT based iPOND dataset and could be divided into 3 groups: enriched in nascent, restored within 11 minutes and delayed (Fig. 3A). As shown previously (Alvarez et al., 2023), TFs defined as pioneers bound newly replicated DNA with similar kinetics as other TFs (Fig. 3B). TFs containing a DNA-binding domain on the other hand tended to bind more transiently to newly replicated chromatin (Fig. 3C). There was no distinction in the mode of binding between TFs involved in activation or repression of transcription (Fig. 3D). This further confirmed that the ability of TFs to regain access to newly replicated DNA benefits from DNA binding ability, not a pioneer status or involvement in transcription activation or repression.

**Figure 3:**
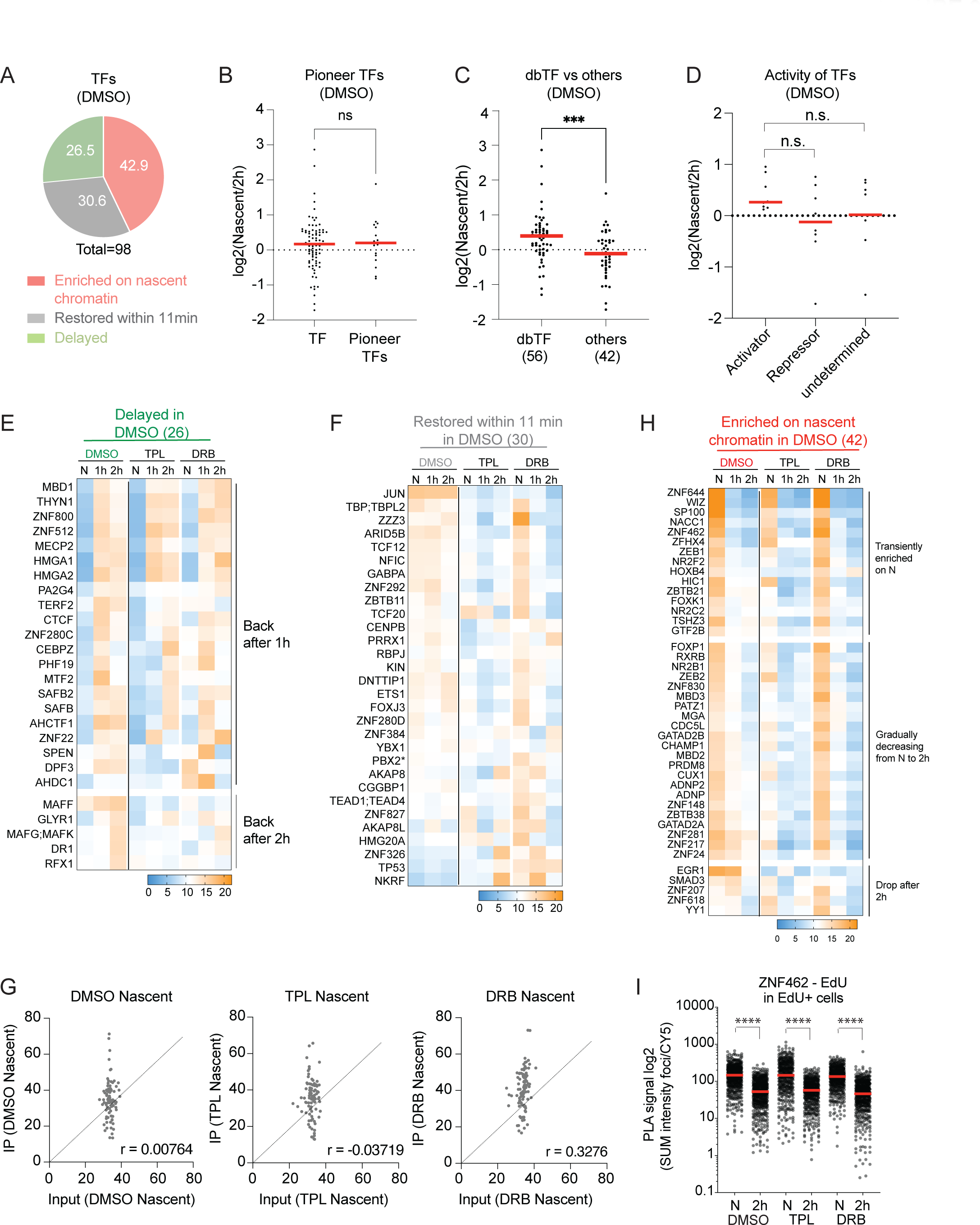
Abundance of TF transiently enriched on nascent chromatin is insensitive to transcription inhibition. **A**. Pie chart showing the number of identified TFs as percentage in three subgroups using criteria defined in Fig. 1I. The list of human TFs was generated based on two studies (Lambert et al., 2018; Lovering et al., 2021). **B**. Graph showing the relative percentage of enrichment on nascent chromatin of pioneer TFs vs. non-Pioneer TFs in DMSO treated cells. Red line: median; p value was calculated using an unpaired t test. **C**. Same as in (B) for TFs classified into two groups, DNA-binding TFs vs others as defined in (Lovering et al., 2021). **D**. Same as in (B) for TFs classified into Activator, Repressor, or undetermined based on (Berest et al., 2019). **E, F**. Heatmap showing the relative enrichment of the two groups defined in DMSO. Each column represents a time point (N: Nascent, 1h, and 2h) and each row corresponds to the protein indicated on the left. The sum of each row corresponds to 100% of enrichment. Colour scale is indicated. The number of TFs in each group is shown on the top. **G**. Scatter plots comparing the relative percentage of enrichment between nascent iPOND samples and the respective input samples in DMSO (*left*), TPL (*middle*), and DRB (*right*) treated cells. The Pearson correlation coefficient (r) is shown. **H.** Heatmap as in E and F, for the TFs defined as enriched in nascent chromatin by DMSO. **I**. Single-cell PLA signal of EdU-ZNF462 interaction shown for nascent chromatin (N) and 2 h mature chromatin (2 h) in DMSO vs. TPL or DRB treated cells. PLA signal was calculated as the SUM of the total intensity of PLA foci per nucleus and normalized according to EdU intensity per cell. >488 nuclei were analysed per sample. Red line, median; Standard deviation is shown. Unpaired and two-sided t test was used. N=3, one representative experiment is shown.

Upon TPL and DRB treatments, TFs from the delayed category, defined by the DMSO condition, retained their ability to bind newly replicated DNA, although in some cases, their reassembly was hurried or further delayed (Fig. 3E). For the TFs from the “restored within 11 min” category, TPL treatment reduced their binding to newly replicated DNA, while DRB treatment increased binding to nascent DNA (Fig. 3F). Therefore, association of this class of TFs with nascent chromatin is consolidated by the presence of RNAPIIS5P. At later time points, there is a reduction in binding compared to DMSO which may indicate a requirement for RNAPII to enable these factors to persist on newly replicated DNA. Of note, although TFs are in general low abundant proteins, there is no correlation between TFs ability to bind newly replicated DNA and their cellular abundance (Fig. 3G, S3A).

Finally, a large proportion of TFs associated with newly replicated chromatin at higher levels than observed for other time points (Fig. 3H). This is consistent with previous observations (Alvarez et al., 2023) and interestingly, TPL or DRB treatments had no impact on their eviction from newly replicated chromatin. This was surprising as sites of spurious chromatin accessibility observed in nascent chromatin over coding regions are to be removed as a result of transcription (Stewart-Morgan et al., 2019). To independently verify our observation, we adopted an approach based on the use of Proximity Ligation Assay (PLA) (Fig. S3B). To do this, ZNF462 was selected as an example of TF highly enriched on nascent chromatin and PLA was used to measure its proximity with EdU labelled nascent DNA. Quantification of the PLA signal using QIBC confirms that ZNF462 is enriched on newly replicated DNA in comparison to mature chromatin (two hours) and that this enrichment is not affected by TPL or DRB treatments (Fig. 3I). Altogether these results reveal that the majority of TFs associate with nascent DNA independently of RNAPII. None the less, the association of approximately 1/3 of the TFs detected in our study (the “restored within 11 min” category) is augmented by the presence of RNAPII.

### Transcription promotes H3.3K36me2 restoration while challenging H3K9me3 incorporation

The last family of proteins which abundance on nascent chromatin was perturbed upon transcription inhibition were proteins involved in histone post translational modifications (Fig. 2B, 2C). This includes histone modifiers involved in imposing both, modifications linked to gene transcription and gene silencing (Fig. S4A), suggesting that following DNA replication, transcription may contribute to histone modification re-establishment (Fig. 4A). To test this hypothesis, we directly monitored the incorporation of modified histone and the restoration of histone modifications on newly replicated chromatin upon transcription inhibition. Histone modifications were detected by label free mass spectrometry from histones isolated by histone extraction (Sidoli et al., 2016) and by iPOND (Fig. 4B). This comparison allowed to assess side by side the level of each modification on total chromatin (histone extraction) and on replicated chromatin (iPOND). The level of each histone modification identified on a residue is expressed as a percentage of all the possible modified states detected for this residue (Table S2). For instance, on total chromatin for the peptides that contains lysine 20 in histone H4, 21% of the peptides are unmodified, 37.8% monomethylated, 40.6% dimethylated, 0.3% trimethylated and 0.14% are acetylated.

**Figure 4:**
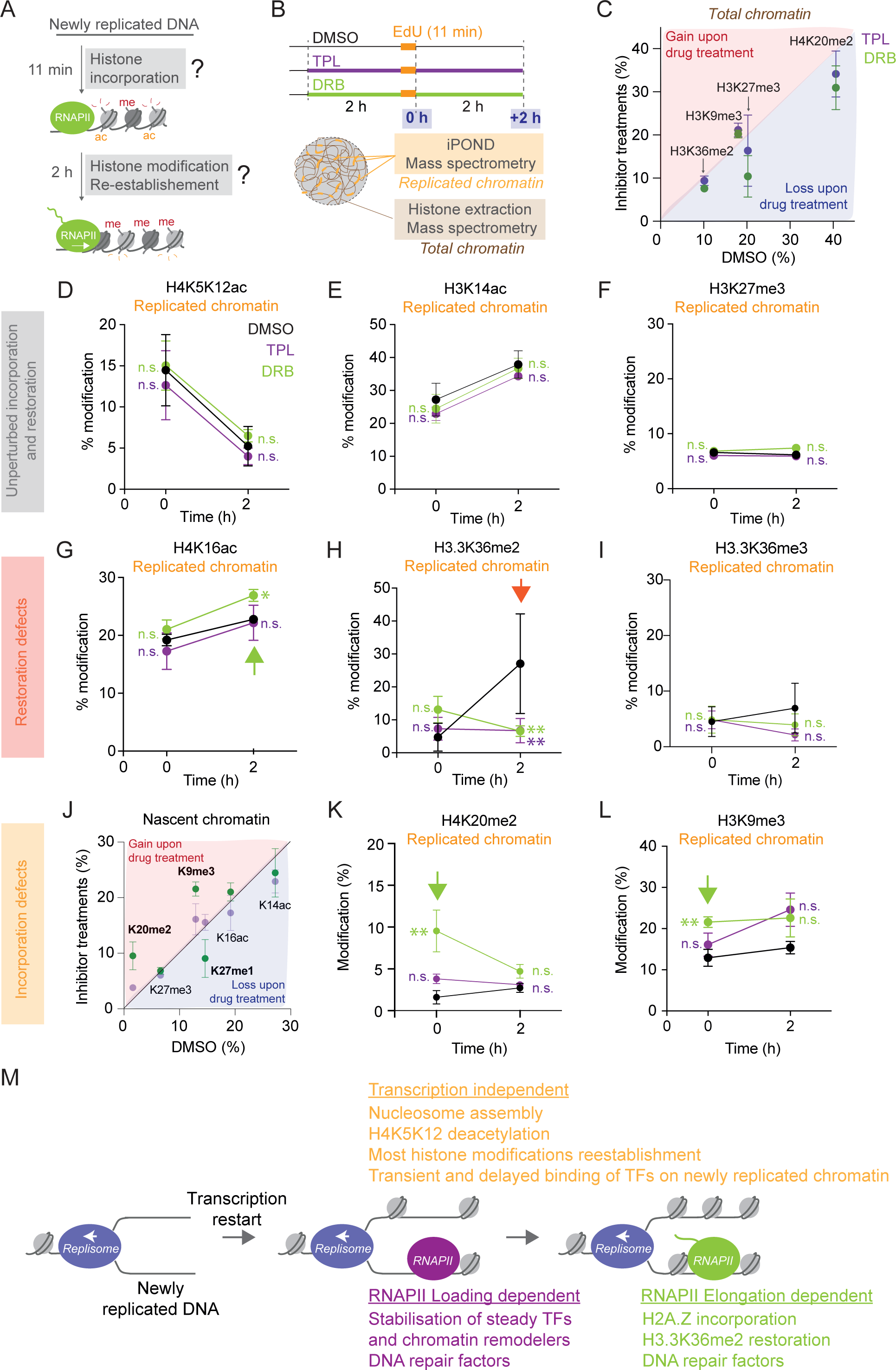
Transcription promotes H.33K36me2 re-establishment. **A**. Behind the replication fork, nascent chromatin is highly acetylated on H4K5K12, and the methylation level are diluted. To which extent transcription contributes to the re-establishment of histone modifications following DNA replication is unknown. **B**. Experimental design of the iPOND-TMT time course experiment in TIG-3 cells in DMSO, TPL or DRB. **C**. Histone modifications on total chromatin after 2 and 4 hours of TPL or DRB treatments. Standard error of the mean is shown. **D**. Proportion of H4K5K12ac on newly replicated chromatin for each time point in DMSO (black), DRB (green) and TPL (purple) treated cells. Standard error of the mean is shown. **E-I**. Same as in (D) for H3K14ac, H3K27me3, H4K16ac, H3.3K36me2 and me3. **J**. Histone modifications on nascent chromatin. Standard error of the mean is shown. **K, L**. Same as in (D) for H4K20me2 and H3K9me3. **M.** Model summarising the key findings from this study. On top is shown the transcription independent events. “Transient binding of TF on newly replicated chromatin” refers to TFs that are enriched on nascent chromatin in DMSO (Fig. 3H). Below, processes and proteins which binding is stabilized or promoted by the binding of RNAPII and RNAPII elongation. “Steady” TFs refers to TFs that are restored within 11 minutes in DMSO (Fig. 3F).

On total chromatin, after DRB or TPL treatments, the level of several histone modifications was perturbed compared to DMSO although none of these changes were significant (Fig. 4C, S4B-K). Of note, the abundance of H3K4me3 was too low to be confidently examined by mass spectrometry (H3K4me3 represents only 0.04 to 0.07% of the H3 3-8 peptide). H3K4me3 on total chromatin was therefore examined by western blot (Fig. S4B), and consistent with transcription-dependent recruitment of H3K4me3-methylating complexes (Zhang et al., 2015), the level of H3K4me3 decreases upon TPL or DRB treatments. On nascent chromatin, we first monitored H4K5 and K12 acetylation, a di-acetylation present mainly on newly synthesized histones and removed within 30 minutes after their incorporation into nucleosome (Benson et al., 2006). Consistent with this, on replicated chromatin, H4K5K12ac levels decrease within two hours, confirming that nascent (0hr) and mature chromatin (2hr) have been successfully isolated (Fig. 4D). Interestingly, neither TPL nor DRB treatments affected the rate of histone H4K5K12 deacetylation. Moreover, at the 0-hr time point, a similar level of diacetylated histone H4 (14%) was detected in DMSO, TPL and DRB treated cells. Taken together this reveals that blocking transcription does not affect the deposition of newly synthesized histones on newly replicated DNA, and does not interfere with the rapid deacetylation of histone H4K5K12.

Similar to H4K5K12 deacetylation (Fig. 4D), restoration of the majority of histone modifications was also unaffected by DRB or TPL treatments (Fig. 4E, 4F, Table S2). The two exceptions were H4K16ac and H3.3K36me2, whose restoration kinetics were significantly affected by DRB or TPL (Fig. 4G, 4H). At the 2-hr time point, H4K16 acetylation was significantly increased in DRB treated cells, while H3.3K36me2 restoration was blocked upon TPL or DRB treatments. Of note, H3.3K36me3 restoration was also affected but not significantly (Fig. 4I). Therefore, in general, transcription does not play a role in the reestablishment of histone modifications following DNA replication, apart from promoting H3.3K36me2 and weakening H4K16ac. Moreover, although transcription inhibition affected reassociation of several histone modifiers (Fig. S4A), this did not translate into changes of their cognate modifications within the time frame of this analysis.

Finally, H4K20me2 and H3K9me3 levels were transiently increased upon DRB treatment (Fig. 4J-L, S4L, S4M, time point 0 h). As di and tri-methylations require several hours to be re-established following DNA replication (Alabert et al., 2015; Reveron-Gomez et al., 2018), 11 minutes after the passage of the fork, such increase most probably reflects an increase of H4K20me2 and H3K9me3 on recycled parental histones. Taken together, these results revealed that on newly replicated DNA, blocking transcription does not affect the deposition of newly synthesized histones. However, blocking transcription elongation leads to transient increase of H4K20me2 and H3K9me3. Once incorporated, histone modification restoration is generally uncoupled from transcription. The two exceptions were the restoration of H3.3K36me2 and H4K16ac.

## DISCUSSION

In this study we analysed the reassembly of chromatin following DNA replication and identified proteins with dependencies on transcription (Fig. 4M). By using two transcription inhibitors, TPL and DRB, we were able to distinguish whether these dependencies are reliant on RNAPII loading or RNAPII elongation. We show that inhibition of RNAPII loading impairs the recruitment of most chromatin remodellers, 1/3 of TFs, H2A.Z, and prevents the reestablishment of normal levels of H3.3K36me2. In addition, blocking RNAPII elongation provokes the transient accumulation of H3K9me3 and H4K20me2, and the gradual accumulation of H1.X. Finally, RNAPII loading and elongation behind replisomes leads to the recruitment of DNA repair factors involved in handling replication transcription conflicts and fork stalling.

We show that on newly replicated DNA, RNAPII binding, but not its elongation, promotes the recruitment of chromatin remodelling complexes (Fig. 2L). Intriguingly, a recent study using similar transcription inhibitors coupled to CUT&TAG showed that in mES cells, RNAPII promotes SMARCA4 binding to chromatin at TSS (Brahma and Henikoff, 2023). Here, RNAPII mediated stabilisation of SMARCA4 is specific to nascent chromatin rather than total chromatin (Fig. 2O). Mechanistically, the stabilisation of SMARCA4 association mediated by RNAPII may explain why active transcription promotes nucleosome positioning at TSS on newly replicated DNA (Stewart-Morgan et al., 2019; Vasseur et al., 2016). However, it is unlikely that repositioning on newly replicated chromatin relies on SMARCA4 alone. Indeed, several chromatin remodellers and histone chaperones have been linked to DNA replication, from playing a role in DNA replication origin activation to replication fork progression (Ahmad et al., 2022; Kurat et al., 2017). Consistent with this, most chromatin remodellers and many histone chaperones were hyper-abundant on newly replicated chromatin in our dataset (Fig. 2D, 2H-L) and in other cell lines (Alvarez et al., 2023). Yet it remains unclear which function each chromatin remodeller has on newly replicated chromatin. The combination of multiple rapid degradation strategies (Blumli et al., 2021; Bomber et al., 2023) will be required to establish functional interplay between chromatin remodellers on newly replicated chromatin.

Our data suggests that on newly replicated DNA, pioneer TFs associate with kinetics similar to other TFs. However, TFs containing a DNA binding domain tend to become transiently enriched on newly replicated DNA. In this study we find that the eviction of these TFs from newly replicated chromatin is not dependent on transcription. The observation of this transient overloading is interesting as it resonates with previous ideas that replication provides a window of opportunity for establishment of new gene regulatory architectures in daughter cells (Stewart-Morgan et al., 2020). It is also consistent with previous observations that sites of spurious chromatin accessibility may arise in newly replicated chromatin over coding genes (Stewart-Morgan et al., 2019). In this case, transcription was observed to restore coding gene chromatin. As the majority of the genome is not transcribed at high frequency alternative mechanisms are likely to regulate where TFs remain stably associated within intergenic chromatin. One possibility is that where TFs bind and can form multivalent interactions, they are more likely to remain associated. Many TFs can form direct or indirect interactions with RNAPII. In this respect RNAPII may act as a hub stabilising via looping interactions the association of TFs with adjacent loci. The effects of transcriptional inhibitors on factor loading are broadly consistent with this. Prevention of RNAPII loading with TPL results in reduced association of a broad range of TFs, chromatin remodelling enzymes and histone modifying enzymes that may be capable of forming direct or indirect interactions with RNAPII. DRB treatment which results in transient enrichment of RNAPII-S5P causes transient enrichment of many of these factors. Obtaining further support for this model may be challenging as the association of many factors in S-phase may be more distributed than at other stages of the cell cycle and as a result more difficult to detect by genomics approaches. Yet, if true, from a cell fate point of view, the access on newly replicated DNA to previously prohibited gene regulatory regions, or the replication coupled “window of opportunity” hypothesis, may be narrower in time than anticipated. Studies in systems with defined transcriptional changes will allow to further explore this process, and to identify the functional differences between the different groups of TFs.

We find that histones are less abundant on nascent chromatin compared to mature chromatin (Fig. 2E). This is consistent with previous studies of newly replicated DNA (Annunziato, 2015), and together with chromatin accessibility studies (Ramachandran and Henikoff, 2016; Stewart-Morgan et al., 2019) it suggests that on nascent chromatin nucleosomes are both, poorly positioned and less abundant. Analysis of modified histone incorporation on newly assembled histones revealed that newly synthesized histones deposition and parental histone recycling is uncoupled from transcription. The two exceptions were H3K9me3 and H4K20me2, as for both modifications, their levels transiently increased on newly replicated chromatin. One possibility could be that the transient increased level of H3K9me3 is the result of DNA damage on newly replicated DNA (Tsouroula et al., 2016). However, we did not detect significant signs of DNA damage in DRB treated cells (Fig. 2A, 2B). Although modelling has predicted that such eventuality alone would not be sufficient to initiate heterochromatin formation (Hathaway et al., 2012), it will be important to further explore this possibility in disease contexts such as cancer, where transcription and replication programs are compromised (Kotsantis et al., 2018). Analysis of the restoration of histone modifications, from 0 hr to 2 hr, revealed that most H3 and H4 histone modifications are removed or re-established independently of transcription. The notable exception is H3.3K36me2 (Fig. 4H). The transcription dependency of H3.3K36me2 is consistent with the fact that H3K36me2 relies, in part, on SET-2 direct binding to RNAPII phosphorylated on S2 and S5 (Kizer et al., 2005; Sun et al., 2005). To our knowledge, H3K4me3 and H2BK120ub1, are the only other known modifications whose restoration on newly replicated chromatin is driven by transcription (Flury et al., 2023; Reveron-Gomez et al., 2018; Serra-Cardona et al., 2022).

Finally, we show that upon transcription inhibition, several repair proteins were depleted from newly replicated chromatin. Because they share the same DNA template, transcription is known to challenge replisome progression at high frequency, from RNAPII constituting a roadblock to progressing replisomes, to generate RNA:DNA hybrids (Berti et al., 2020). It is therefore remarkable that behind replisomes, only a handful of DNA repair factors appear to be involved in response to RNAPII binding and elongation. One interesting factor is SMARCAL1 which is recruited to replisomes in DMSO and DRB treated cells. As SMARCAL1 promotes fork reversal allowing replication fork to restart (Berti et al., 2020; Betous et al., 2012), it argues in favour of a model where in each S phase, fork reversal is required to deal with the high frequency of transcription replication conflicts. Finally, using a RNAPII degron, a recent study has elegantly demonstrated that transcription upon exit from mitosis promotes the reorganization of high order structures in G1 phase (Zhang et al., 2021). In our system, CTCF (Fig. 3E), nucleopores (Table S1) and cohesins (Fig. S1G) were perturbed by DRB and TPL treatments but only moderately. Therefore, a causative relationship between transcription and nuclear organization behind replisomes remains to be established. Altogether this study reveals that RNAPII promotes association of a diverse selection of factors with nascent chromatin. It will be interesting to investigate whether transcription has additional roles during differentiation when programmed transcriptional changes must be implemented.

### Limitations of the study

The proteomics approach we have taken is not locus specific meaning that we cannot distinguish the association of factors to transcriptionally active or inactive regions. As a small proportion of the genome is transcriptionally active, it is possible our study underestimates transcription dependencies that only occur at transcribed regions. On the other hand, it enables to avoid cell cycle synchronization that is known to introduce changes on chromatin (Ly et al., 2015). Furthermore, it remains unknown how much of the changes detected on replicated chromatin would be applicable to total chromatin. Proteomic profiling of chromatin fractions after transcription inhibition has been performed but in different cell lines and using harsher transcription inhibition conditions (Skalska et al., 2021). Therefore, it will be interesting to repeat these experiments in our system to identify processes specific to newly replicated chromatin. Finally, the usage of transcription inhibitors could be avoided with the recent development of RNAPII degron cell lines, allowing to explore the role of RNAPII on newly replicated chromatin with greater precision.

## ACKNOWLEDGEMENTS

We thank Giulia Saredi, Kathleen Stewart-Morgan and Jane Wright for reading the manuscript. We thank the imaging facilities in School of Life Sciences. S.B. is supported by the ERC-Stg-IDRE. V.A. is supported by the CRUK-CDF C57404/A21782. H.J. was supported by Wellcome Collaborator Award (Ref: 206293/Z/17/Z). Research in the Owen-Hughes lab is funded by MRC grant MR/SO21647/1. The Sidoli lab gratefully acknowledges for funding the Einstein-Mount Sinai Diabetes center, Merck, Relay Therapeutics, Deerfield (Xseed award) and the NIH Office of the Director (S10OD030286). Research in the Alabert lab is funded by CRUK-CDF (C57404/A21782) and the European Research Council ERC-Stg-IDRE.

## AUTHOR CONTRIBUTIONS

Conceptualization, S.B. and C.A.; Methodology, S.B. and C.A; Validation V.A. and S.B; Formal Analysis, S.B., V.A., H.J., S.G., S.S., and C.A.; Investigation, S.B., V.A., H.J., S.G., R.S., M.T., S.S., and C.A.; Writing – Original Draft, S.B. and C.A.; Writing – Review & Editing, all the authors; Funding Acquisition, C.A.

## MATERIALS AND METHODS

### Cell culture

The normal fibroblast cell line TIG-3-20, derived from Japanese fetal lung (JCRB0506, JCRB Cell Bank) was cultured in Dulbecco’s Modified Eagle’s Medium (DMEM, Gibco, 61965059) supplemented with 10% FBS fetal bovine serum (FBS, Gibco, 10270106) and 1% penicillin/streptomycin at 37°C in 5% CO_2_ and humid environment. For all large-scale iPOND experiments, 4 x 10^6^ cells were seeded in 15 cm dishes 2 days prior to EdU labeling and processing, 10 dishes per sample.

### DNA Labelling

All samples were labeled with medium containing 5-ethynyl-20-deoxyuridine (EdU; Thermo Fisher scientific, E10187) at a final concentration of 20 μM for 11 minutes. Following labeling, nascent samples were immediately taken and processed. All mature samples were washed twice in 1X PBS and further incubated in fresh media containing 20 μM thymidine (Sigma, T1895) for the appropriate time interval before collection.

### Transcription Inhibition

To inhibit transcription initiation, 2 hours prior EdU labeling, media was changed for media containing triptolide (Sigma, T3652) at a final concentration of 1 μM. To block transcription elongation, 2 hours prior EdU labeling, media was changed for media containing 5,6-dichlorobenzimida-zole 1-b-D-ribofuranoside (DRB; Sigma, D1916) at a final concentration of 50 μM. DMSO-treated controls were treated 2 hours prior to EdU labeling with 10 μl DMSO/plate (Sigma, D8418). Inhibitors and DMSO were included for the duration of EdU labeling and chromatin maturation in all experiments.

### Isolation of proteins in nascent DNA (iPOND)

iPOND was performed as described previously (Sirbu et al., 2012). In short, 1 × 10^8^ asynchronised TIG-3-20 cells (per time point) were labeled with 20 μM EdU (Thermo Fisher) for 11 min. The thymidine chased sample were washed twice and subsequently further incubated in medium containing 20 μM thymidine for the indicated times. All samples were crosslinked with 1% formaldehyde (Sigma, F8775) for 15 min at RT followed by 5 min incubation with 0.125 M glycine to quench the formaldehyde. Afterward, cells were scraped from the plate (on ice), permeabilized in 0.25% Triton X-100 in PBS for 30 min at RT and washed with 0.5% BSA in PBS. To conjugate biotin to EdU-labeled DNA, click chemistry reactions were performed for 1.5 h in the dark (10 μM Biotin-azide [Thermo Fisher, B10184], 10 mM Sodium ascorbate, 2 mM CuSO_4_ in 1xPBS). Cells were lysed (1% SDS in 50 mM Tris, pH 8.0, protease inhibitor), sonicated (Bioruptor, Diagenode), the lysate diluted 1:1 (v/v) with cold PBS containing protease inhibitor, and streptavidin beads (Thermo Fisher, 65002) used to capture the biotin-conjugated DNA-protein complexes. At this point, the protein lysate is collected to be analyzed together with the iPOND pulldown. Captured complexes were washed extensively using lysis buffer and 1 M NaCl. The proteins were eluted under reducing conditions by boiling in 2X LSB sample buffer for 30 min.

### Tandem mass tag (TMT)-based MS for quantitative proteomics analysis (TMT MS)

iPOND samples were prepared using SP3 protocol for TMT labeling as described in (Hughes et al., 2014). In brief, protein samples were mixed with SP3 beads (1: 10, protein: beads) and digested with trypsin (1: 50, trypsin: protein). For total extracts, equal amounts (100 μg) of each peptide sample were dried and dissolved in 100 μL 100 mM TEAB buffer (pH 8.0). For iPOND samples, all the peptides from each sample were dried and dissolved in 100 μL 100 mM TEAB buffer (pH 8.0). TMT labeling of each sample were followed by the TMT10plex Isobaric Mass Tag Labeling Kit (Thermo Fisher Scientific) manual. The TMT labeled peptide samples were further fractionated using offline high-pH reverse-phase (RP) chromatography, as previously described (Brenes et al., 2019). The 24 fractions were subsequently dried and the peptides re-dissolved in 5% formic acid and analyzed by nanoLC–MS/MS system Orbitrap Fusion Tribrid Mass Spectrometer (Thermo Fisher Scientific), equipped with an UltiMate 3000 RSLCnano Liquid Chromatography System, as previously described (Brenes et al., 2019) The MS data were analyzed using MaxQuant (v 1.6.7.0) and searched against Homo Sapiens database from Uniport (Swissport, downloaded at January 2020) (Cox and Mann, 2011). The TMT quantification was set to reporter ion MS3 type with 10plex TMT (LOT: UH285228). The detailed parameter file was uploaded together with the raw MS data. Protein groups output table from MaxQuant was cleaned, filtered and median normalised with Perseus (1.6.7.0) (Tyanova et al., 2016). ***Mass spectrometry data analysis.*** MaxQuant software was used for the identification and TMT quantification of proteins. The datasets were analyzed and filtered with Perseus. Proteins that were more abundant in the negative control (no EdU labeling, + ClickIT) compared to the 2 h DMSO mature sample were removed from all time points since considered as background/non-specific. Proteins identified as ‘Potential contaminants’, ‘Only identified by site’, or ‘Reverse’ were removed and only proteins that are identified in 3/3 experiments as well as have one or more unique peptides were kept in the analysis. The datasets were median normalised, and the percentage of enrichment calculated in Microsoft Excel. Heatmaps, scatter plots, correlation heatmaps, Principal Component Analysis (PCA) as well as volcano plots were generated in GraphPrism. R was used to perform hierarchical clustering (k = 22), script available upon request. Gene Ontology analysis of biological process as well as Kyoto Encyclopedia of Genes and Genomes (KEGG) was performed using STRING and g:Profiler.

### Immunoblotting

Total cell extracts were prepared using 2X Laemmli Sample buffer (100mM Tris-HCl (pH 6.8), 4% SDS, Glycerol) and the protein amount was determined using the Pierce BCA Protein Assay (23227, Thermo Fisher Scientific). After measuring the protein concentration, NuPAGE® Sample reducing agents (NP0009, Invitrogen) and NuPAGE® LDS Sample Buffer (NP0007, Invitrogen) were added to the samples. After boiling at 95°C for 5 min, the cell extracts or de-crosslinked samples were subjected to SDS–PAGE separation on NuPAGE 4–12% gels (Invitrogen, NP0321) in MES buffer. Proteins were transferred onto 0.2 μm pore size nitrocellulose membranes (GE Healthcare, 10600001) at 18 V for 1 h using Trans-Blot SD Semi-Dry Transfer Cell (Bio-Rad). Membranes were blocked in 5% skimmed milk in tris-buffer saline (TBS, 50 mM Tris, pH 7.5, 150 mM NaCl) supplemented with 0.1% Tween20 (TBS-T) for 1 h. The following dilution for each antibody were used: Anti-PCNA (1:1000, Abcam, ab29), anti-RNA polymerase II CTD repeat YSPTSPS [phospho S5] [4H8] (1:1000, ab5408, Abcam), anti-histone H3 (1:1000, ab10799, Abcam), anti-H3K4me3 (1:1000, ab8580, Abcam), anti-GAPDH (1:2000, 2118, Cell Signalling Technology), rabbit-HRP (1:5000, 115-035-062, Jackson Immunoresearch), and mouse-HRP (1:5000, 711-035-152, Jackson Immunoresearch). Signals from HRP-conjugated antibodies were revealed by SuperSignal™ West Pico PLUS Chemiluminescent (Thermo Fisher Scientific, 34580). Membranes were imaged using ChemiDoc XRS+ (Bio-Rad).

### Microscopy

*Immunofluorescence.* TIG-3-20 cells were seeded at 10 000 cells per well in clear bottom, black 96-well plates (Corning, 3340) and grown for 24 h. To measure DNA replication and transcription rates, cells were treated either with 20 μM EdU for 20 min or 1 mM 5-ethynyluridine (EU) for 1h, respectively, prior performing pre-extraction with ice-cold CSK buffer (10 mM PIPES (pH 7), 100 mM NaCl, 300 mM Sucrose, 3 mM MgCl_2_). After pre-extraction cells were wash once with 1X PBS and fixed with 2% formaldehyde for 20 min. For EdU detection, Alexa 647 was covalently linked to EdU using the Click-iT EdU Imaging Kit (Thermo Fisher Scientific, C10640). For EU detection, Alexa 488 was covalently linked to EU using the Click-iT RNA Alexa Flour 488 Imaging Kit (Thermo Fisher Scientific, C10329). Samples were blocked with BSA in PBS-Tween and antibodies were incubated at the following dilutions: anti-Biotin (1:1000, Vector Labs, SP-3000), anti-RNA polymerase II CTD repeat YSPTSPS [phospho S5] [4H8] (1:1000, ab5408, Abcam), anti-RNA polymerase II CTD repeat YSPTSPS [8WG16] (1:500, ab817, Abcam), anti-SMARCA4 (1:500, ab110641, Abcam), anti-rabbit IgG AF488 (1:1000, 1910751, Invitrogen), anti-Alexa Fluor 546 goat anti-mouse IgG(H+L) (1:1000, A11030, Invitrogen), and anti-Alexa Fluor 488 donkey anti-mouse IgG(H+L) (1:1000, A21202, Invitrogen). For DNA staining, the secondary antibodies were incubated together with DAPI (Thermo Fisher Scientific, 62248). The cells were then washed twice in PBS-Tween, one time with PBS, and left in PBS until imaging. *Proximal ligation assay (PLA).* Cells were fixed and permeabilised as described before. EdU was then covalently linked to biotin-azide (Thermo Fisher Scientific, B10184) using the Click-iT EdU Imaging Kit (Thermo Fisher Scientific, C10340). To allow data normalisation using the EdU signal, the Click-iT reaction was performed in the presents of Alexa647-azide that was spiked in using a 40x dilution to the Biotin-azide. PLA between biotin and the protein of interest was performed according to manufacturer instructions, using the Duolink® Proximity Ligation Assay from Sigma (DUO92006, DUO92014, DUO92002) or the Proximity Ligation kit from Navinci (Cambridge bioscience, NF.GR.100). After performing the PLA protocol, cells were stained with DAPI (Thermo Fisher Scientific, 62248) for 20 min in PBS, washed with PBS, and left in PBS until imaging. *QIBC.* Images were taken and analyzed with ScanR High Content Screening Microscopy (Olympus). Data were visualised and statistically analyzed in Tableau and GraphPrism.

### Remodeler Assay *in vitro*

Nucleosomes were reconstituted on Cy3 labelled DNA, based on the 601 sequence, with a 17bp DNA extension on one side and a 47 bp extension on the other side of the 601 sequence. Repositioning by Brg1 subcomplex was performed in 20 mM HEPES pH7.5, 50 mM NaCl, 3 mM MgCl_2_, 1 mM ATP, 100 nM each nucleosome, and 20 nM Brg1 in a 20 uL reaction for each condition with specified amount of either DMSO, TPL or DRB. The reactions were carried out on ice and stopped with the addition of 100 ng/µL competitor DNA, 120 mM NaCl, and 2% sucrose. Repositioned nucleosomes were run on 6% PAGE/0.2X TBE gels in recirculating 0.2X TBE buffer for 3–4 hr at 250V. The percent of repositioned nucleosomes was analysed using Aida image analysis software and plotted in Microsoft Excel.

### Measuring PTMs on newly replicated chromatin upon transcription inhibition

The iPOND was performed as described before with two changes. The captured complexes were washed with lysis buffer and 1 M NaCl, followed by two washes with 50 mM Tris, pH 8. The cells for the total protein lysate were taken before crosslinking (approx. 8×10^6^ cells) and processed using the histone extraction protocol. ***Histone extraction and digestion***. TIG3 cells were washed twice with PBS and the cell pellets then resuspended in ice cold Triton Extraction Buffer (TEB: PBS containing 0.5% Triton X-100, AEBSF, Protease Inhibitor Cocktail) at a cell density of 10e7 cells per ml. After lysing on ice for 10 min, cells were centrifuged at 2000 rpm for 10 min at 4°C and supernatant was discarded. To remove the residual cytoplasmic protein, the cell pellets were washed one more time with the TEB buffer and centrifuged again as before. The cell pellets were resuspended in 0.2 N HCl at a proper density (4×10e7 cells/ml) and vortexed overnight at 4°C. The supernatant acid extracts were collected after 10 minutes centrifugation at 6500 rpm. Equal volume of 50% trichloroacetic acid (TCA) was added and mixed on ice for 30 min. After centrifugation for 10 min at 10,000 rpm the pellets were washed with ice-cold acetone and centrifuged again at 10,000 rpm for 10 min. The samples were dried in a vacuum centrifuge. The pellet was dissolved in 50 mM ammonium bicarbonate, pH 8.0, and histones were subjected to derivatization using 5 µL of propionic anhydride and 14 µL of ammonium hydroxide (all Sigma Aldrich) to balance the pH at 8.0. The mixture was incubated for 15 min and the procedure was repeated. Histones were then digested with 1 µg of sequencing grade trypsin (Promega) diluted in 50mM ammonium bicarbonate (1:20, enzyme:sample) overnight at room temperature. Derivatization reaction was repeated to derivatize peptide N-termini. The samples were dried in a vacuum centrifuge. ***Sample desalting.*** Prior to mass spectrometry analysis, samples were desalted using a 96-well plate filter (Orochem) packed with 1 mg of Oasis HLB C-18 resin (Waters). Briefly, the samples were resuspended in 100 µl of 0.1% TFA and loaded onto the HLB resin, which was previously equilibrated using 100 µl of the same buffer. After washing with 100 µl of 0.1% TFA, the samples were eluted with a buffer containing 70 µl of 60% acetonitrile and 0.1% TFA and then dried in a vacuum centrifuge. ***LC-MS/MS Acquisition and Analysis*** Samples were resuspended in 10 µl of 0.1% TFA and loaded onto a Dionex RSLC Ultimate 300 (Thermo Scientific), coupled online with an Orbitrap Fusion Lumos (Thermo Scientific). Chromatographic separation was performed with a two-column system, consisting of a C-18 trap cartridge (300 µm ID, 5 mm length) and a picofrit analytical column (75 µm ID, 25 cm length) packed in-house with reversed-phase Repro-Sil Pur C18-AQ 3 µm resin. Peptides were separated using a 30 min gradient from 1-30% buffer B (buffer A: 0.1% formic acid, buffer B: 80% acetonitrile + 0.1% formic acid) at a flow rate of 300 nl/min. The mass spectrometer was set to acquire spectra in a data-independent acquisition (DIA) mode. Briefly, the full MS scan was set to 300-1100 m/z in the orbitrap with a resolution of 120,000 (at 200 m/z) and an AGC target of 5×10e5. MS/MS was performed in the orbitrap with sequential isolation windows of 50 m/z with an AGC target of 2×10e5 and an HCD collision energy of 30. Histone peptides raw files were imported into EpiProfile 2.0 software (Sidoli et al., 2016). From the extracted ion chromatogram, the area under the curve was obtained and used to estimate the abundance of each peptide. In order to achieve the relative abundance of post-translational modifications (PTMs), the sum of all different modified forms of a histone peptide was considered as 100% and the area of the particular peptide was divided by the total area for that histone peptide in all of its modified forms. The relative ratio of two isobaric forms was estimated by averaging the ratio for each fragment ion with different mass between the two species. The resulting peptide lists generated by EpiProfile were exported to Microsoft Excel and further processed for a detailed analysis.

### Quantification and statistical analysis

For figure 1B-E, 2M-O, 3B-D, unpaired t tests were performed in GraphPrism. For Figure 2L and 4D-L, paired t tests were performed in GraphPrism. For PLA data, all t tests applied were unpaired and two-sided using GraphPrism. Pearson correlations were performed with a confidence interval of 95% and two-tailed in GraphPrism. Principal Components Analysis were performed in GraphPrism with a 95% of percentile level and 1000 simulations.

## Data availability

The TMT mass spectrometry proteomics data have been deposited to the ProteomeXchange Consortium via the PRIDE (Perez-Riverol et al., 2022) partner repository with the dataset identifier PXD040888 and are publicly available as of the date of publication. Mass spectrometry raw files of the label free analysis of the histone modifications are available on the public repository Chorus (https://chorusproject.org) under the project number 1812. Any additional information required to reanalyse the data reported in this paper is available from the lead contact upon request.

## References

1. Ahmad, K., Henikoff, S., and Ramachandran, S. (2022). Managing the Steady State Chromatin Landscape by Nucleosome Dynamics. Annu Rev Biochem 91, 183–195.

2. Alabert, C., Barth, T.K., Reveron-Gomez, N., Sidoli, S., Schmidt, A., Jensen, O.N., Imhof, A., and Groth, A. (2015). Two distinct modes for propagation of histone PTMs across the cell cycle. Genes Dev 29, 585–590.

3. Alvarez, V., Bandau, S., Jiang, H., Rios-Szwed, D., Hukelmann, J., Garcia-Wilson, E., Wiechens, N., Griesser, E., Ten Have, S., Owen-Hughes, T., et al. (2023). Proteomic profiling reveals distinct phases to the restoration of chromatin following DNA replication. Cell Rep 42, 111996.

4. Annunziato, A.T. (2015). The Fork in the Road: Histone Partitioning During DNA Replication. Genes (Basel) 6, 353–371.

5. Benson, L.J., Gu, Y., Yakovleva, T., Tong, K., Barrows, C., Strack, C.L., Cook, R.G., Mizzen, C.A., and Annunziato, A.T. (2006). Modifications of H3 and H4 during chromatin replication, nucleosome assembly, and histone exchange. J Biol Chem 281, 9287–9296.

6. Berest, I., Arnold, C., Reyes-Palomares, A., Palla, G., Rasmussen, K.D., Giles, H., Bruch, P.M., Huber, W., Dietrich, S., Helin, K., et al. (2019). Quantification of Differential Transcription Factor Activity and Multiomics-Based Classification into Activators and Repressors: diffTF. Cell Rep 29, 3147–3159 e3112.

7. Berti, M., Cortez, D., and Lopes, M. (2020). The plasticity of DNA replication forks in response to clinically relevant genotoxic stress. Nat Rev Mol Cell Biol 21, 633–651.

8. Betous, R., Mason, A.C., Rambo, R.P., Bansbach, C.E., Badu-Nkansah, A., Sirbu, B.M., Eichman, B.F., and Cortez, D. (2012). SMARCAL1 catalyzes fork regression and Holliday junction migration to maintain genome stability during DNA replication. Genes Dev 26, 151–162.

9. Blumli, S., Wiechens, N., Wu, M.Y., Singh, V., Gierlinski, M., Schweikert, G., Gilbert, N., Naughton, C., Sundaramoorthy, R., Varghese, J., et al. (2021). Acute depletion of the ARID1A subunit of SWI/SNF complexes reveals distinct pathways for activation and repression of transcription. Cell Rep 37, 109943.

10. Bomber, M.L., Wang, J., Liu, Q., Barnett, K.R., Layden, H.M., Hodges, E., Stengel, K.R., and Hiebert, S.W. (2023). Human SMARCA5 is continuously required to maintain nucleosome spacing. Mol Cell 83, 507–522 e506.

11. Brahma, S., and Henikoff, S. (2023). RNA Polymerase II, the BAF remodeler and transcription factors synergize to evict nucleosomes. bioRxiv.

12. Brenes, A., Hukelmann, J., Bensaddek, D., and Lamond, A.I. (2019). Multibatch TMT Reveals False Positives, Batch Effects and Missing Values. Mol Cell Proteomics 18, 1967–1980.

13. Cox, J., and Mann, M. (2011). Quantitative, high-resolution proteomics for data-driven systems biology. Annu Rev Biochem 80, 273–299.

14. Dobrinic, P., Szczurek, A.T., and Klose, R.J. (2021). PRC1 drives Polycomb-mediated gene repression by controlling transcription initiation and burst frequency. Nat Struct Mol Biol 28, 811–824.

15. Ebmeier, C.C., Erickson, B., Allen, B.L., Allen, M.A., Kim, H., Fong, N., Jacobsen, J.R., Liang, K., Shilatifard, A., Dowell, R.D., et al. (2017). Human TFIIH Kinase CDK7 Regulates Transcription-Associated Chromatin Modifications. Cell Rep 20, 1173–1186.

16. Flury, V., Reveron-Gomez, N., Alcaraz, N., Stewart-Morgan, K.R., Wenger, A., Klose, R.J., and Groth, A. (2023). Recycling of modified H2A-H2B provides short-term memory of chromatin states. Cell 186, 1050–1065 e1019.

17. Giaimo, B.D., Ferrante, F., Herchenrother, A., Hake, S.B., and Borggrefe, T. (2019). The histone variant H2A.Z in gene regulation. Epigenetics Chromatin 12, 37.

18. Hathaway, N.A., Bell, O., Hodges, C., Miller, E.L., Neel, D.S., and Crabtree, G.R. (2012). Dynamics and memory of heterochromatin in living cells. Cell 149, 1447–1460.

19. Hughes, C.S., Foehr, S., Garfield, D.A., Furlong, E.E., Steinmetz, L.M., and Krijgsveld, J. (2014). Ultrasensitive proteome analysis using paramagnetic bead technology. Mol Syst Biol 10, 757.

20. Iurlaro, M., Stadler, M.B., Masoni, F., Jagani, Z., Galli, G.G., and Schubeler, D. (2021). Mammalian SWI/SNF continuously restores local accessibility to chromatin. Nat Genet 53, 279-287.

21. Kim, J.M., Visanpattanasin, P., Jou, V., Liu, S., Tang, X., Zheng, Q., Li, K.Y., Snedeker, J., Lavis, L.D., Lionnet, T., et al. (2021). Single-molecule imaging of chromatin remodelers reveals role of ATPase in promoting fast kinetics of target search and dissociation from chromatin. Elife 10.

22. Kizer, K.O., Phatnani, H.P., Shibata, Y., Hall, H., Greenleaf, A.L., and Strahl, B.D. (2005). A novel domain in Set2 mediates RNA polymerase II interaction and couples histone H3 K36 methylation with transcript elongation. Mol Cell Biol 25, 3305–3316.

23. Kotsantis, P., Petermann, E., and Boulton, S.J. (2018). Mechanisms of Oncogene-Induced Replication Stress: Jigsaw Falling into Place. Cancer Discov 8, 537–555.

24. Kurat, C.F., Yeeles, J.T.P., Patel, H., Early, A., and Diffley, J.F.X. (2017). Chromatin Controls DNA Replication Origin Selection, Lagging-Strand Synthesis, and Replication Fork Rates. Mol Cell 65, 117–130.

25. Lambert, S.A., Jolma, A., Campitelli, L.F., Das, P.K., Yin, Y., Albu, M., Chen, X., Taipale, J., Hughes, T.R., and Weirauch, M.T. (2018). The Human Transcription Factors. Cell 175, 598–599.

26. Li, J., Cai, Z., Vaites, L.P., Shen, N., Mitchell, D.C., Huttlin, E.L., Paulo, J.A., Harry, B.L., and Gygi, S.P. (2021). Proteome-wide mapping of short-lived proteins in human cells. Mol Cell 81, 4722–4735 e4725.

27. Lovering, R.C., Gaudet, P., Acencio, M.L., Ignatchenko, A., Jolma, A., Fornes, O., Kuiper, M., Kulakovskiy, I.V., Laegreid, A., Martin, M.J., et al. (2021). A GO catalogue of human DNA-binding transcription factors. Biochim Biophys Acta Gene Regul Mech 1864, 194765.

28. Ly, T., Endo, A., and Lamond, A.I. (2015). Proteomic analysis of the response to cell cycle arrests in human myeloid leukemia cells. Elife 4.

29. Millan-Arino, L., Izquierdo-Bouldstridge, A., and Jordan, A. (2016). Specificities and genomic distribution of somatic mammalian histone H1 subtypes. Biochim Biophys Acta 1859, 510–519.

30. Neri, F., Rapelli, S., Krepelova, A., Incarnato, D., Parlato, C., Basile, G., Maldotti, M., Anselmi, F., and Oliviero, S. (2017). Intragenic DNA methylation prevents spurious transcription initiation. Nature 543, 72–77.

31. Owens, N., Papadopoulou, T., Festuccia, N., Tachtsidi, A., Gonzalez, I., Dubois, A., Vandormael-Pournin, S., Nora, E.P., Bruneau, B.G., Cohen-Tannoudji, M., et al. (2019). CTCF confers local nucleosome resiliency after DNA replication and during mitosis. Elife 8.

32. Padeken, J., Methot, S.P., and Gasser, S.M. (2022). Establishment of H3K9-methylated heterochromatin and its functions in tissue differentiation and maintenance. Nat Rev Mol Cell Biol 23, 623–640.

33. Palozola, K.C., Donahue, G., Liu, H., Grant, G.R., Becker, J.S., Cote, A., Yu, H., Raj, A., and Zaret, K.S. (2017). Mitotic transcription and waves of gene reactivation during mitotic exit. Science 358, 119–122.

34. Perez-Riverol, Y., Bai, J., Bandla, C., Garcia-Seisdedos, D., Hewapathirana, S., Kamatchinathan, S., Kundu, D.J., Prakash, A., Frericks-Zipper, A., Eisenacher, M., et al. (2022). The PRIDE database resources in 2022: a hub for mass spectrometry-based proteomics evidences. Nucleic Acids Res 50, D543–D552.

35. Pommier, Y., Nussenzweig, A., Takeda, S., and Austin, C. (2022). Human topoisomerases and their roles in genome stability and organization. Nat Rev Mol Cell Biol 23, 407–427.

36. Ramachandran, S., and Henikoff, S. (2016). Transcriptional Regulators Compete with Nucleosomes Post-replication. Cell 165, 580–592.

37. Reveron-Gomez, N., Gonzalez-Aguilera, C., Stewart-Morgan, K.R., Petryk, N., Flury, V., Graziano, S., Johansen, J.V., Jakobsen, J.S., Alabert, C., and Groth, A. (2018). Accurate Recycling of Parental Histones Reproduces the Histone Modification Landscape during DNA Replication. Mol Cell 72, 239–249 e235.

38. Schick, S., Grosche, S., Kohl, K.E., Drpic, D., Jaeger, M.G., Marella, N.C., Imrichova, H., Lin, J.G., Hofstatter, G., Schuster, M., et al. (2021). Acute BAF perturbation causes immediate changes in chromatin accessibility. Nat Genet 53, 269–278.

39. Serra-Cardona, A., Duan, S., Yu, C., and Zhang, Z. (2022). H3K4me3 recognition by the COMPASS complex facilitates the restoration of this histone mark following DNA replication. Sci Adv 8, eabm6246.

40. Sidoli, S., Bhanu, N.V., Karch, K.R., Wang, X., and Garcia, B.A. (2016). Complete Workflow for Analysis of Histone Post-translational Modifications Using Bottom-up Mass Spectrometry: From Histone Extraction to Data Analysis. J Vis Exp.

41. Sirbu, B.M., Couch, F.B., and Cortez, D. (2012). Monitoring the spatiotemporal dynamics of proteins at replication forks and in assembled chromatin using isolation of proteins on nascent DNA. Nat Protoc 7, 594–605.

42. Sirbu, B.M., Couch, F.B., Feigerle, J.T., Bhaskara, S., Hiebert, S.W., and Cortez, D. (2011). Analysis of protein dynamics at active, stalled, and collapsed replication forks. Genes Dev 25, 1320–1327.

43. Skalska, L., Begley, V., Beltran, M., Lukauskas, S., Khandelwal, G., Faull, P., Bhamra, A., Tavares, M., Wellman, R., Tvardovskiy, A., et al. (2021). Nascent RNA antagonizes the interaction of a set of regulatory proteins with chromatin. Mol Cell 81, 2944–2959 e2910.

44. Steurer, B., Janssens, R.C., Geverts, B., Geijer, M.E., Wienholz, F., Theil, A.F., Chang, J., Dealy, S., Pothof, J., van Cappellen, W.A., et al. (2018). Live-cell analysis of endogenous GFP-RPB1 uncovers rapid turnover of initiating and promoter-paused RNA Polymerase II. Proc Natl Acad Sci U S A 115, E4368–E4376.

45. Stewart-Morgan, K.R., Petryk, N., and Groth, A. (2020). Chromatin replication and epigenetic cell memory. Nat Cell Biol 22, 361–371.

46. Stewart-Morgan, K.R., Reveron-Gomez, N., and Groth, A. (2019). Transcription Restart Establishes Chromatin Accessibility after DNA Replication. Mol Cell 75, 284–297 e286.

47. Sun, X.J., Wei, J., Wu, X.Y., Hu, M., Wang, L., Wang, H.H., Zhang, Q.H., Chen, S.J., Huang, Q.H., and Chen, Z. (2005). Identification and characterization of a novel human histone H3 lysine 36-specific methyltransferase. J Biol Chem 280, 35261–35271.

48. Toledo, L.I., Altmeyer, M., Rask, M.B., Lukas, C., Larsen, D.H., Povlsen, L.K., Bekker-Jensen, S., Mailand, N., Bartek, J., and Lukas, J. (2013). ATR prohibits replication catastrophe by preventing global exhaustion of RPA. Cell 155, 1088–1103.

49. Tsouroula, K., Furst, A., Rogier, M., Heyer, V., Maglott-Roth, A., Ferrand, A., Reina-San-Martin, B., and Soutoglou, E. (2016). Temporal and Spatial Uncoupling of DNA Double Strand Break Repair Pathways within Mammalian Heterochromatin. Mol Cell 63, 293–305.

50. Tyanova, S., Temu, T., Sinitcyn, P., Carlson, A., Hein, M.Y., Geiger, T., Mann, M., and Cox, J. (2016). The Perseus computational platform for comprehensive analysis of (prote)omics data. Nat Methods 13, 731–740.

51. Vasseur, P., Tonazzini, S., Ziane, R., Camasses, A., Rando, O.J., and Radman-Livaja, M. (2016). Dynamics of Nucleosome Positioning Maturation following Genomic Replication. Cell Rep 16, 2651–2665.

52. Venkatesh, S., Smolle, M., Li, H., Gogol, M.M., Saint, M., Kumar, S., Natarajan, K., and Workman, J.L. (2012). Set2 methylation of histone H3 lysine 36 suppresses histone exchange on transcribed genes. Nature 489, 452–455.

53. Vispe, S., DeVries, L., Creancier, L., Besse, J., Breand, S., Hobson, D.J., Svejstrup, J.Q., Annereau, J.P., Cussac, D., Dumontet, C., et al. (2009). Triptolide is an inhibitor of RNA polymerase I and II-dependent transcription leading predominantly to down-regulation of short-lived mRNA. Mol Cancer Ther 8, 2780–2790.

54. Zhang, S., Ubelmesser, N., Josipovic, N., Forte, G., Slotman, J.A., Chiang, M., Gothe, H.J., Gusmao, E.G., Becker, C., Altmuller, J., et al. (2021). RNA polymerase II is required for spatial chromatin reorganization following exit from mitosis. Sci Adv 7, eabg8205.

55. Zhang, T., Cooper, S., and Brockdorff, N. (2015). The interplay of histone modifications - writers that read. EMBO Rep 16, 1467–1481.

