## Supplemental Figure 1-4 for "Transcription promotes the restoration of chromatin following DNA replication"

### FIGURE S1

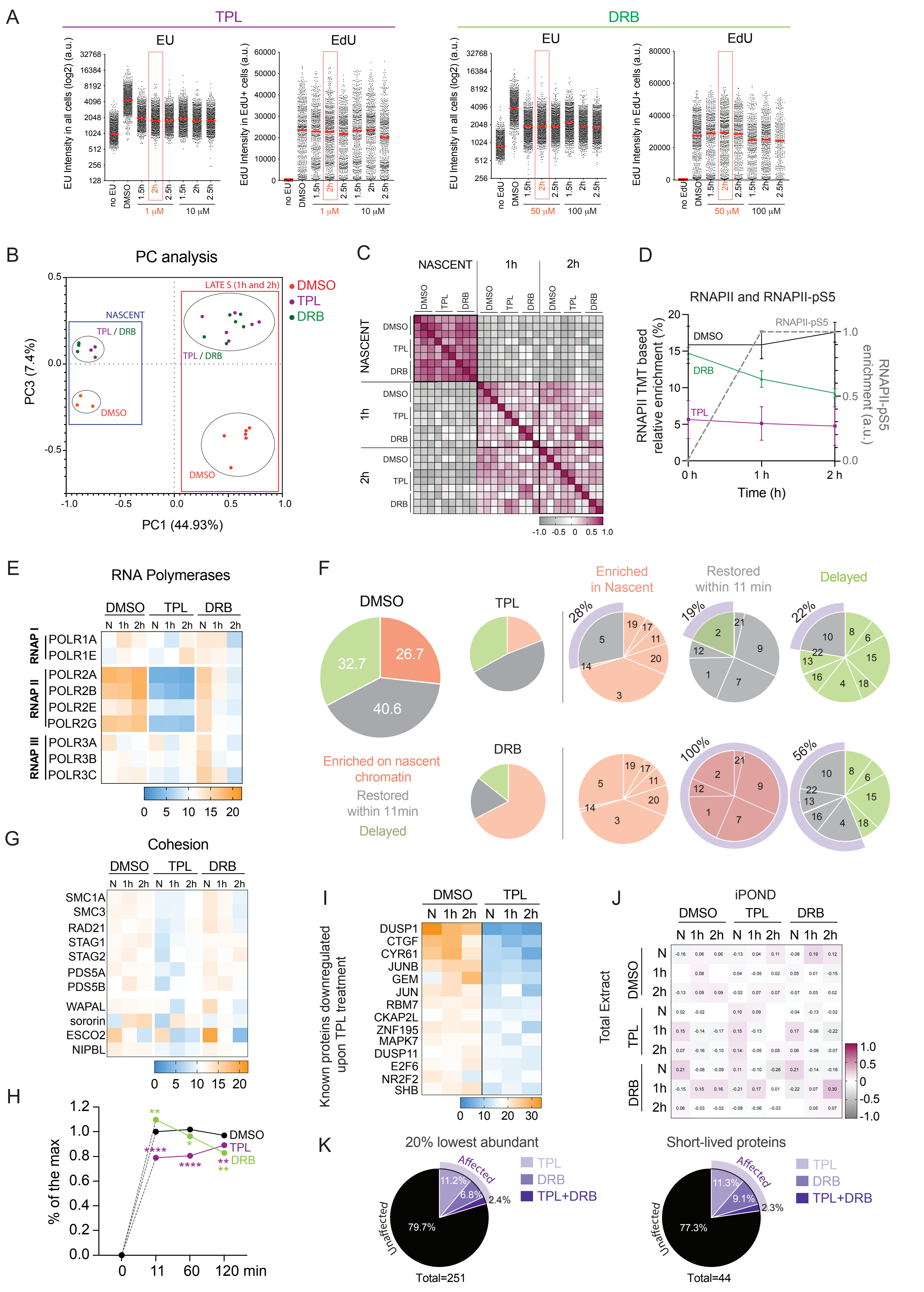
A
TPL
DRB
EU
EdU
EU Intensity in all cells (log2) (a.u.)
EdU Intensity in EdU+ cells (a.u.)
no EU
no EdU
DMSO
1.5h
2h
2.5h
1 μM
10 μM
50 μM
100 μM
B
PC analysis
NASCENT
TPL / DRB
DMSO
LATE S (1h and 2h)
PC3 (7.4%)
PC1 (44.93%)
C
NASCENT
1h
2h
D
RNAPII and RNAPII-pS5
RNAPII TMT based relative enrichment (%)
RNAPII-pS5 enrichment (a.u.)
Time (h)
E
RNA Polymerases
RNAP I
RNAP II
RNAP III
POLR1A
POLR1E
POLR2A
POLR2B
POLR2E
POLR2G
POLR3A
POLR3B
POLR3C
N 1h 2h
F
Enriched in Nascent
Restored within 11 min
Delayed
Enriched on nascent chromatin
Restored within 11min
28%
19%
22%
100%
56%
32.7
26.7
40.6
G
Cohesion
SMC1A
SMC3
RAD21
STAG1
STAG2
PDS5A
PDS5B
WAPAL
sororin
ESCO2
NIPBL
H
% of the max
120 min
I
Known proteins downregulated upon TPL treatment
DUSP1
CTGF
CYR61
JUNB
GEM
JUN
RBM7
CKAP2L
ZNF195
MAPK7
DUSP11
E2F6
NR2F2
SHB
J
iPOND
Total Extract
K
20% lowest abundant
Short-lived proteins
Affected
Unaffected
TPL+DRB
11.2%
6.8%
2.4%
79.7%
Total=251
11.3%
9.1%
2.3%
77.3%
Total=44

SUPPLEMENTARY FIGURE LEGENDS

**Figure S1: Proteomic profiling of chromatin behind replisomes upon transcription inhibition. Related to Figure 1**. **A**. 5-ethynyluridine (EU) and 5-Ethynyl-2'-deoxyuridine (EdU) incorporation measured upon transcription inhibition with TPL or DRB by QIBC. For EU, the log2 intensity of all cells is shown, for EdU, the intensity of EdU positive cells is shown for each time point and concentration tested. Concentration and timing chosen for the large scale iPOND-TMT experiment is highlighted in orange. **B**. Principal-component analysis of the three biological replicates based on the proteins commonly identified. **C.** Pearson correlation heatmap between time points of the three biological replicates. Colour scale is indicated. **D.** Average of the relative enrichment of RNAPII subunits between time points in DMSO (black), TPL (purple), and DRB (green) treated cells. The grey dotted line represents the enrichment of RNAPII-pS5 from (Fig. 1H). **E.** Heatmaps of relative enrichment of RNAPI, II, and III (n = 3). Each column represents a time point (N: Nascent, 1h, and 2h) and each row corresponds to the protein indicated on the left. The sum of each row corresponds to 100% of enrichment. Colour scale is indicated. **F.** Pie chart of protein percentage from the hierarchical clustering shown in Fig 1I. **G**. Same as in (E) for Cohesin proteins. **H**. Quantification of the Cohesin related proteins. N=3, statistic based on a paired t test. **I.** Same as in (E) for known proteins downregulated upon TPL treatment based on (Vispe et al., 2009). Shown is the relative enrichment between DMSO and TPL using their abundance in total extracts. **J.** Pearson correlation heatmap between iPOND and total extract samples using the average of relative enrichment of the three biological replicates. Colour scale is indicated. **K.** Pie chart showing the effect of transcription inhibition as measured in figure 1J, on the 20% lowest abundant proteins identified by iPOND (left), and the 44 proteins with the shortest half-life (right).

### Figure S2

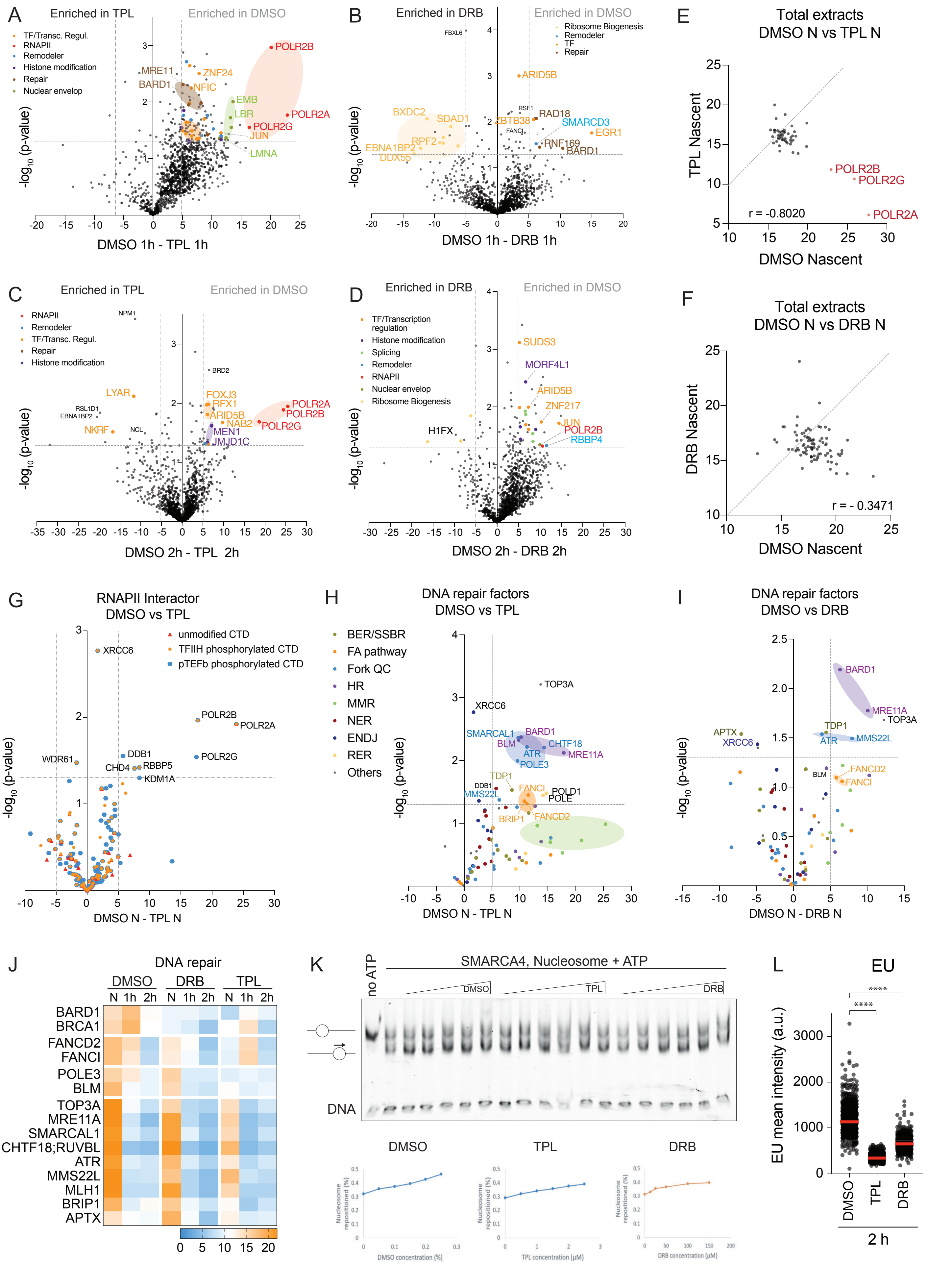
A
Enriched in TPL
Enriched in DMSO
TF/Transc. Regul.
RNAPII
Remodeler
Histone modification
Repair
Nuclear envelop
POLR2B
MRE11
BARD1
ZNF24
NFIC
EMB
LBR
POLR2A
POLR2G
JUN
LMNA
-log10 (p-value)
DMSO 1h - TPL 1h
B
Enriched in DRB
Enriched in DMSO
Ribosome Biogenesis
Remodeler
TF
Repair
FBXL6
ARID5B
BXDC2
SDAD1
RSF1
RAD18
ZBTB38
SMARCD3
FANCI
EGR1
RPF2
RNF169
EBNA1BP2
BARD1
DDX55
DMSO 1h - DRB 1h
E
Total extracts
DMSO N vs TPL N
TPL Nascent
POLR2B
POLR2G
r = -0.8020
POLR2A
DMSO Nascent
C
Enriched in TPL
Enriched in DMSO
RNAPII
Remodeler
TF/Transc. Regul.
Repair
Histone modification
NPM1
BRD2
LYAR
FOXJ3
RFX1
POLR2A
POLR2B
RSL1D1
EBNA1BP2
ARID5B
NAB2
POLR2G
NKRF
NCL
MEN1
JMJD1C
DMSO 2h - TPL 2h
D
Enriched in DRB
Enriched in DMSO
TF/Transcription regulation
Histone modification
Splicing
Remodeler
RNAPII
Nuclear envelop
Ribosome Biogenesis
SUDS3
MORF4L1
ARID5B
ZNF217
H1FX
JUN
POLR2B
RBBP4
DMSO 2h - DRB 2h
F
Total extracts
DMSO N vs DRB N
DRB Nascent
r = - 0.3471
DMSO Nascent
G
RNAPII Interactor
DMSO vs TPL
unmodified CTD
TFIIH phosphorylated CTD
pTEFb phosphorylated CTD
XRCC6
POLR2B
POLR2A
WDR61
DDB1
CHD4
RBBP5
POLR2G
KDM1A
DMSO N - TPL N
H
DNA repair factors
DMSO vs TPL
BER/SSBR
FA pathway
Fork QC
HR
MMR
NER
ENDJ
RER
Others
TOP3A
XRCC6
SMARCAL1
BARD1
BLM
CHTF18
ATR
MRE11A
POLE3
TDP1
DDB1
FANCI
POLD1
MMS22L
POLE
BRIP1
FANCD2
DMSO N - TPL N
I
DNA repair factors
DMSO vs DRB
BARD1
MRE11A
TOP3A
APTX
TDP1
XRCC6
ATR
MMS22L
FANCD2
BLM
FANCI
DMSO N - DRB N
J
DNA repair
DMSO
DRB
TPL
N 1h 2h N 1h 2h N 1h 2h
BARD1
BRCA1
FANCD2
FANCI
POLE3
BLM
TOP3A
MRE11A
SMARCAL1
CHTF18;RUVBL
ATR
MMS22L
MLH1
BRIP1
APTX
0 5 10 15 20
K
no ATP
SMARCA4, Nucleosome + ATP
DMSO
TPL
DRB
DNA
DMSO
TPL
DRB
Nucleosome repositioned (%)
DMSO concentration (%)
TPL concentration (μM)
DRB concentration (μM)
L
EU
EU mean intensity (a.u.)
DMSO
TPL
DRB
2 h
****
****

**Figure S2: Chromatin remodellers abundance on nascent and steady state chromatin is impaired upon TPL treatment. Related to figure 2. A-D.** Enrichment of proteins across DRB and TPL iPOND-TMT-MS datasets. **E, F.** Scatter plots comparing the relative percentage of enrichment between cellular extracts in DMSO, TPL and DRB treated cells. The Pearson correlation coefficient (r) is shown. **G.** Enrichment of unmodified RNAPII CTD, TFIIF phosphorylated RNAPII CTD and pTEFb phosphorylated RNAPII CTD interactors. **H, I.** Enrichment of DNA repair proteins across DRB and TPL iPOND-TMT-MS datasets. **J.** Relative enrichment of DNA repair proteins identified in Fig. 2A, 2B. **K.** Nucleosome sliding *in vitro* assay (top) and its quantification (bottom). **L**. QIBC analysis of EU level in DMSO, TPL or DRB treated cells. Graphs show the mean intensity per nuclei. Red line: median; p value calculated using an unpaired t test. N=3, one representative experiment is shown.

### Figure S3

A

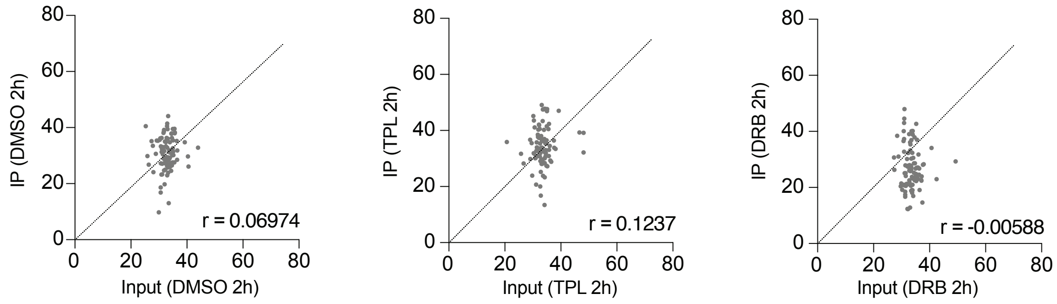

DMSO 2h
IP (DMSO 2h)
Input (DMSO 2h)
r = 0.06974
TPL 2h
IP (TPL 2h)
Input (TPL 2h)
r = 0.1237
DRB 2h
IP (DRB 2h)
Input (DRB 2h)
r = -0.00588
0
20
40
60
80


B

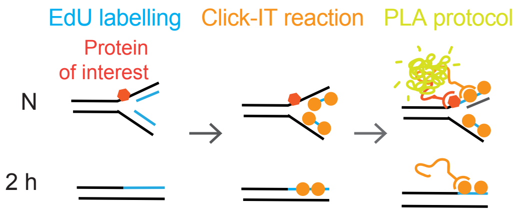

EdU labelling
Click-IT reaction
PLA protocol
Protein
of interest
N
2 h

**Figure S3: Majority of TFs transiently enriched on nascent chromatin is insensitive to transcription inhibition. Related to figure 3. A**. Enrichment between 2h iPOND samples and the respective total cell extract samples in DMSO (*left*), TPL (*middle*), and DRB (*right*) treated cells. The Pearson correlation coefficient (r) is shown. **B**. Scheme of the PLA analysis between EdU and protein of interest by QIBC. TIG-3 cells were EdU labeled, and nascent chromatin and 2 h chase samples were collected. Click-IT chemistry and PLA protocols were performed, and acquired images analysed for DAPI, EdU, and PLA intensities (see details in material and method section)

### FIGURE S4

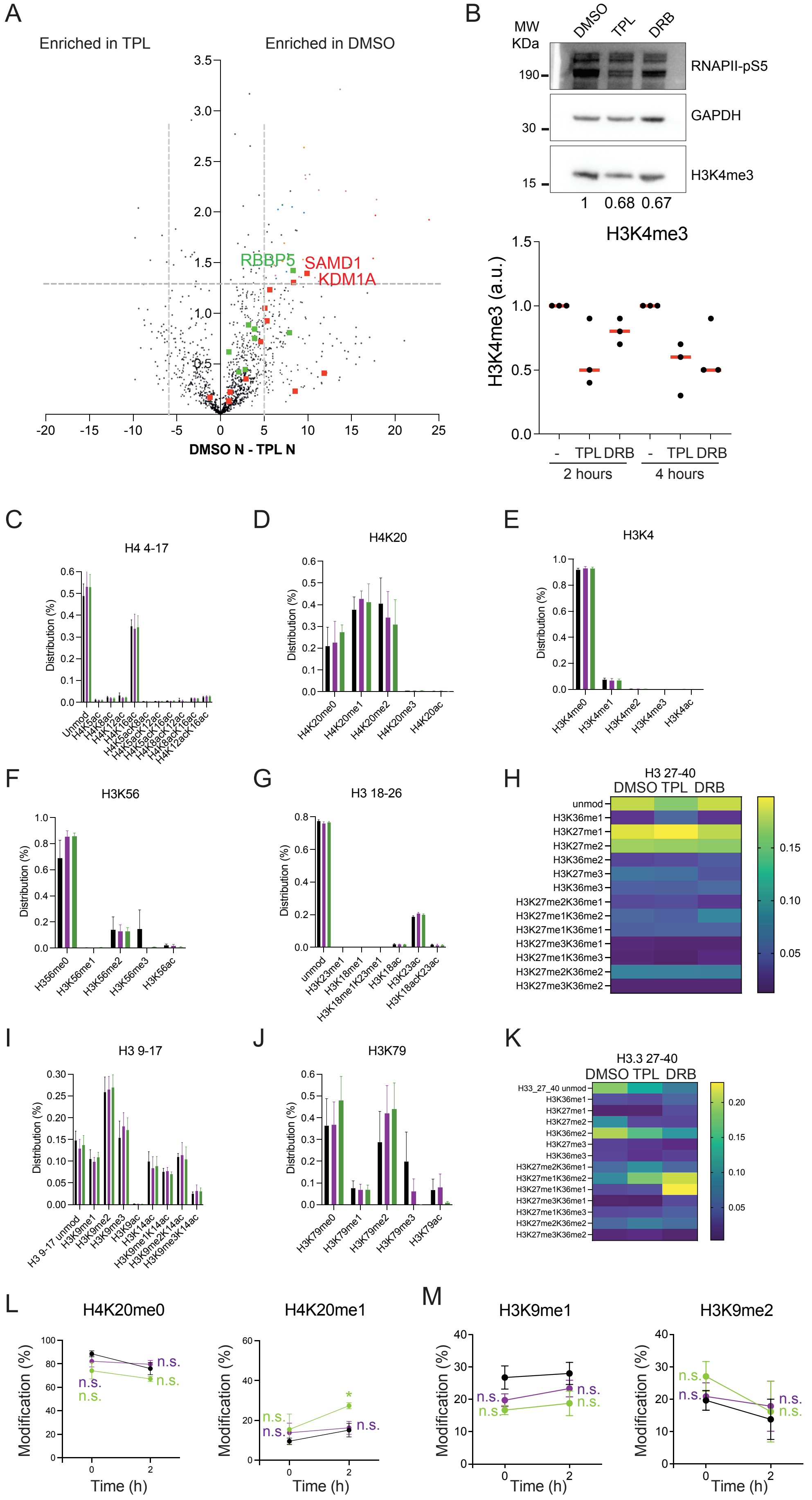
A
Enriched in TPL
Enriched in DMSO
RBBP5
SAMD1
KDM1A
DMSO N - TPL N
B
MW KDa
DMSO TPL DRB
RNAPII-pS5
GAPDH
H3K4me3
190
30
15
1 0.68 0.67
H3K4me3
H3K4me3 (a.u.)
- TPL DRB - TPL DRB
2 hours
4 hours
C
H4 4-17
Distribution (%)
D
H4K20
E
H3K4
F
H3K56
G
H3 18-26
H
H3 27-40
DMSO TPL DRB
I
H3 9-17
J
H3K79
K
H3.3 27-40
DMSO TPL DRB
L
H4K20me0
H4K20me1
M
H3K9me1
H3K9me2
Modification (%)
Time (h)
n.s.

**Figure S4: Transcription restart promotes H3K36me2 re-establishment. Related to figure 4. A.** Enrichment of proteins across TPL iPOND-TMT-MS datasets. Histone modifiers and accessory factors are highlighted. Red, factors linked to transcriptional repression; Green, factors linked to gene activation. Full dataset available in Table S1. **B**. Western Blot analysis of H3K4me3 in total cell extracts. The Western Blot was probed with antibodies against RNAPII-pS5, GABDH, and H3K4me3 (indicated on the right). Sample name indicated on the top. Protein marker indicated on the left. The quantification of the H3K4me3 bands after normalisation using GABDH is shown below (n = 3). **C-K**. Quantification of H3 and H4 modifications from total chromatin extracts. Full dataset available on Table S2. **L, M**. Quantification of H3 and H4 modifications from newly replicated chromatin. Full dataset available on Table S2.

**Table S1**. Proteins identified in 3/3 experiments by iPOND in TIG-3 cells. The average percentages as well as the percentages of each replicate are shown. The cluster number for each protein established based on hierarchical clustering (k = 22), their levels in the proteome sample (average percentage, n = 3) as well as the iBAQ total from the Input sample for each protein are indicated.

**Table S2.** Histone modifications identified in 3 experiments by iPOND (nascent) and histone acid extraction (total chromatin) in TIG3-cells. The average percentages and standard deviation are shown.

References

Vispe, S., DeVries, L., Creancier, L., Besse, J., Breand, S., Hobson, D.J., Svejstrup, J.Q., Annereau, J.P., Cussac, D., Dumontet, C., *et al.* (2009). Triptolide is an inhibitor of RNA polymerase I and II-dependent transcription leading predominantly to down-regulation of short-lived mRNA. Mol Cancer Ther *8*, 2780-2790.
